# From gHBfix to NBfix: Reweighting-Driven Refinement of Hydrogen-Bond Interactions in RNA Force Fields

**DOI:** 10.64898/2026.03.20.713292

**Authors:** Vojtěch Mlýnský, Petra Kührová, Giovanni Bussi, Michal Otyepka, Jiří Šponer, Pavel Banáš

## Abstract

Understanding RNA structural dynamics is essential for elucidating its biological functions, and molecular dynamics (MD) simulations provide an important atomistic complement to experimental approaches. However, the predictive power of MD is fundamentally limited by the accuracy of the underlying empirical Force Fields (FFs), particularly in capturing the delicate balance of non-bonded interactions. Here, we present a systematic reparameterization strategy that replaces the external gHBfix19 hydrogen-bond (H-bond) correction potential with an equivalent set of NBfix Lennard–Jones modifications within a state-of-the-art RNA FF. Using a quantitatively converged temperature replica-exchange MD ensemble of the GAGA tetraloop, we employed a reweighting-based optimization protocol to derive NBfix parameters that reproduce the thermodynamic effects of the original gHBfix19 terms. Sequential optimization of individual gHBfix19 components proved essential to ensure stable and transferable parameter refinement. The resulting fully reformulated NBfix-based variant, termed OL3_CP_–NBfix19, was validated on a representative set of RNA motifs, including tetranucleotides, A-form duplexes, and tetraloops. Across all tested systems, its performance is comparable to that of the reference gHBfix19 FF. By embedding the H-bond corrections directly into the standard non-bonded framework, the NBfix formulation eliminates external biasing potentials, simplifies practical deployment, and reduces computational overhead. Beyond this specific reparameterization, our results demonstrate a practical workflow for translating targeted H-bond corrections into native FF terms for efficient biomolecular simulations.

## INTRODUCTION

Molecular dynamics (MD) simulations have become an essential tool for studying the structure, dynamics, and folding of nucleic acids at atomic resolution.^1–6^ Over the past decades, continuous improvements in simulation algorithms, enhanced sampling techniques, and computational resources have substantially expanded the range of RNA systems accessible to MD simulations.^4, 7^ At the same time, the reliability and predictive power of these simulations remain critically dependent on the accuracy of the underlying Force Fields (FFs).^8–16^ In particular, even subtle imbalances in the FF parameterization can lead to systematic errors in RNA conformational ensembles and folding thermodynamics, limiting the transferability of simulation results across different RNA motifs and environments.^11, 15, 17–19^

State-of-the-art RNA FFs derived within the AMBER framework have reached a level where major artifacts observed in early simulations, such as irreversible backbone distortions or ladder-like unfolding of A-RNA helices, have largely been eliminated.^8, 20^ However, extensive simulation studies have shown that current state-of-the-art RNA FFs are limited not by obvious deficiencies in canonical helices, but rather by more subtle imbalances in non-bonded interactions that govern the thermodynamic stability and rugged free-energy landscapes of diverse RNA motifs.^11, 15, 17, 19, 21–27^ Importantly, the manifestation of these imbalances is often system-dependent and may involve interdependent effects, complicating their identification and correction.^4, 16, 23, 28, 29^ As a result, accurately reproducing the thermodynamic balance among competing conformational states remains challenging, particularly for benchmark systems such as short oligonucleotides and compact RNA motifs. To address these limitations, several targeted FF corrections have been proposed and systematically evaluated, aiming to selectively rebalance specific interactions while preserving the overall consistency of the parametrization.

Among them, the gHBfix approach introduced an additional, physically transparent correction acting directly on selected classes of hydrogen-bonds (H-bonds).^11^ By strengthening under-stabilized base–base –NH···N– interactions and weakening overly favorable sugar–phosphate –OH···O–contacts, gHBfix was shown to significantly improve the stability of native RNA motifs and the folding behavior of challenging systems such as GNRA tetraloops (TLs).^11, 24^ Subsequent studies further demonstrated that gHBfix provides a flexible framework for fine-tuning H-bond interactions and can be systematically optimized using data-driven or semi-automated machine-learning approaches.^19^ Despite its conceptual clarity and demonstrated effectiveness, the gHBfix formalism also introduces practical limitations. The correction is implemented through additional distance-dependent potential terms that must be explicitly defined for each relevant donor–acceptor pair. This increases the complexity of simulation setup, complicates FF distribution, and introduces a non-negligible computational overhead that scales with the number of corrected interactions. While these costs are acceptable in FF development and benchmarking studies, they limit the routine use of gHBfix-augmented FFs in large-scale simulations or high-throughput applications.

A natural strategy to overcome these limitations is to transfer the physical effects encoded in gHBfix corrections into standard non-bonded FF terms that are natively supported by MD engines. Within the framework of the AMBER FF, this can be achieved using so-called NBfix modifications, which allow pair-specific adjustments of Lennard–Jones (LJ) parameters outside the conventional Lorentz–Berthelot combination rules.^30^ In contrast to gHBfix, NBfix terms are straightforward to implement, do not require additional restraints, and incur no extra computational cost. However, a direct translation of gHBfix corrections into NBfix parameters is non-trivial, as it must preserve the delicate thermodynamic balance achieved by the original H-bond corrections without introducing new artifacts.

In this work, we present a systematic reparameterization of the gHBfix19 potential into an equivalent set of NBfix terms. Building on a quantitatively converged temperature replica-exchange MD simulation (T-REMD)^31^ of the r(gcGAGAgc) TL (GAGA TL), we employ a reweighting-based strategy^32, 33^ to identify optimal LJ parameter modifications that reproduce the thermodynamic effects of individual gHBfix terms. By explicitly monitoring the statistical reliability of the reweighted ensembles, we ensure that the resulting NBfix parameters remain within the domain of validity of the reference FF. The derived NBfix parametrization fully replaces the original gHBfix19 corrections^11^ while preserving the folding equilibrium of the reference GAGA TL. Beyond reweighting, we validate the new NBfix FF variant through extensive T-REMD simulations, first by substituting individual gHBfix terms with their NBfix counterparts and subsequently by replacing the entire gHBfix19 potential. Finally, we assess the basic performance of the resulting FF, denoted as OL3_CP_–NBfix19, on a diverse set of benchmark RNA systems including five tetranucleotides (TNs), two A-RNA duplexes, and two TLs. Together, these results demonstrate that the physical effects of gHBfix can be robustly transferred into a computationally efficient NBfix-based formulation, providing a practical and production-ready RNA FF while retaining the benefits of targeted H-bond corrections.

## METHODS

### Reweighting

Reweighting^34^ was applied to evaluate the thermodynamic consequences of removing each gHBfix19 term and substituting it with an NBfix-modified LJ interaction, allowing us to screen candidate parametrizations without rerunning simulations. All reweighting calculations were based on the equilibrated 298.996 K replica from the reference 25 μs-long T-REMD simulation of the GAGA TL initiated from the folded state, which was published in our recent study.^35^ These simulations were performed with AMBER *ff*99bsc0χ_OL3_ RNA FF (i.e., OL3),^8, 20, 36, 37^ further adjusted by the phosphate oxygen van der Waals correction (CP),^38^ in combination with our reparameterization of the affected α, γ, δ and ζ backbone torsions^9^ and gHBfix19 H-bond correction,^11^ henceforth denoted as OL3_CP_–gHBfix19. The final 5 μs of this simulation were used as the training ensemble.

For each tested NBfix parameter set, the change in non-bonded energy was evaluated as the difference between the reference FF description (standard LJ interactions supplemented by gHBfix19 terms) and the target description in which the corresponding gHBfix19 corrections are removed and replaced by NBfix-modified LJ interactions. Formally, the non-bonded energy difference for a configuration *x_i_* was computed as:

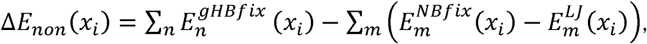

where the first term represents the total contribution of the gHBfix19 potentials acting on all relevant H-bond interactions *n*, and the second term corresponds to the difference between the NBfix-modified LJ interaction and the original LJ interaction for each affected atom-type pair *m*. The gHBfix energy terms were calculated as defined in our previous study,^11^ while the LJ and NBfix interaction energies were evaluated using the standard 12–6 LJ functional form with either the original Lorentz–Berthelot parameters or the modified NBfix parameters, respectively.

The gHBfix19 potential comprises three energy terms: (i) an attractive correction that stabilizes base–base H-bonds by strengthening –NH···N– interactions between amino or imino H and N acceptors (AMBER H, NB and NC atom types) by 1.0 kcal/mol, (ii) a repulsive correction that destabilizes sugar-phosphate H-bonds by weakening –OH···nbO– interactions between ribose 2’-OH hydrogens and phosphate nonbridging oxygens (AMBER HO and OP atom types) by −0.5 kcal/mol, and (iii) an analogous repulsive correction applied to sugar-phosphate H-bonds between between ribose 2’-OH hydrogens and phosphate bridging oxygens (–OH···bO– interactions; AMBER HO and OR atom types), also by −0.5 kcal/mol.

The statistical weight of *i*-th frame in the reweighted ensemble is then:

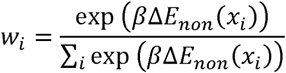

with β = 1⁄*k_B_T*. Any observable *(A)* in the modified FF can be obtained by *(A) = ∑_i_ w_i_A_i_*. In our procedure we estimated reweighted fraction of folded state at ∼298 K that was defined as a total weight of the snapshot having the *ε*RMSD^39^ difference from the native state conformation below 0.7. This quantity served as the primary target observable for matching the behavior of the OL3_CP_–gHBfix19 reference ensemble. Because reweighting becomes unreliable when the target FF deviates too strongly from the control FF used to generate the training ensemble, the individual gHBfix19 terms were reparameterized separately. To quantify the effective statistical support of the reweighted ensemble, we employed the normalized exponential Shannon entropy, hereafter referred to as the normalized effective sample size *P*_eff_:

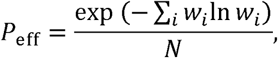

where *N* is number of snapshots in the training set. The *P*_eff_ values close to 1 indicate that a large fraction of the reference ensemble contributes meaningfully to the reweighted distribution, while values approaching zero signal that only a few frames dominate and the reweighting results are therefore unreliable.

### Starting structures and simulation setup

All reweighting calculations and validation simulations were based on the r(gcGAGAgc) 8-mer RNA TL (GAGA TL), which served as the primary reference system. Reference native state (i.e., the folded structure) was taken from the 1.04 Å resolution X-ray structure of the sarcin/ricin loop (PDB ID 1Q9A,^40^ residues 2658-2663) and capped by one additional GC base-pair yielding the final r(gcGAGAgc) sequence. The equilibrated training ensemble used for reweighting was extracted from the final 5 μs of a previously published 25 μs-long T-REMD simulation initiated from the folded state and performed using the OL3_CP_–gHBfix19 FF.^35^ This ensemble corresponds to the 298.996 K replica and was shown to be fully converged with respect to folding–unfolding equilibrium.^35^ All subsequent validation by T-REMD simulations in the present work were started directly from the final snapshots of this reference simulation, ensuring continuity between the training ensemble and newly generated trajectories.

To evaluate the general performance of the derived NBfix-based FF (hereafter denoted OL3_CP_–NBfix19) beyond the reference TL, additional standard MD simulations were carried out on a diverse set of RNA systems, including five TNs, two A-RNA duplexes, and two TL motifs (Table 1). The TN set comprised r(AAAA), r(CAAU), r(CCCC), r(GACC), and r(UUUU) sequences constructed using the Nucleic Acid Builder module of AmberTools18.^41^ The RNA duplex systems included (i) decamer with the r(GCACCGUUGG)_2_ sequence, excised from the crystal structure (PDB ID 1QC0;^42^ 1QC0 duplex), and (ii) r(UUAUAUAUAUAUAA)_2_ tetradecamer (PDB ID 1RNA;^43^ 1RNA duplex). In addition, two TL systems were examined: (i) r(ggcacUUCGgugcc) 14-mer TL (UUCG TL) taken from the NMR ensemble (PDB ID 2KOC,^44^ structure #13), and (ii) the GAGA TL prepared from the 1Q9A^40^ X-ray structure as described above.

**Table 1:**
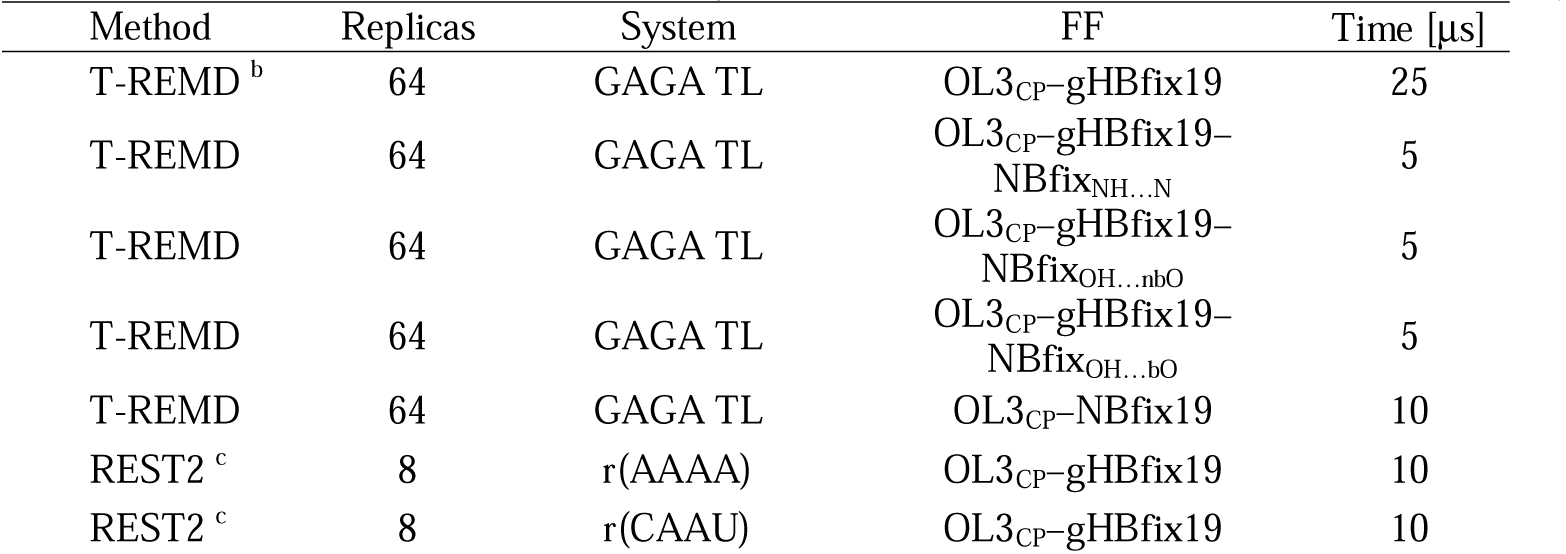
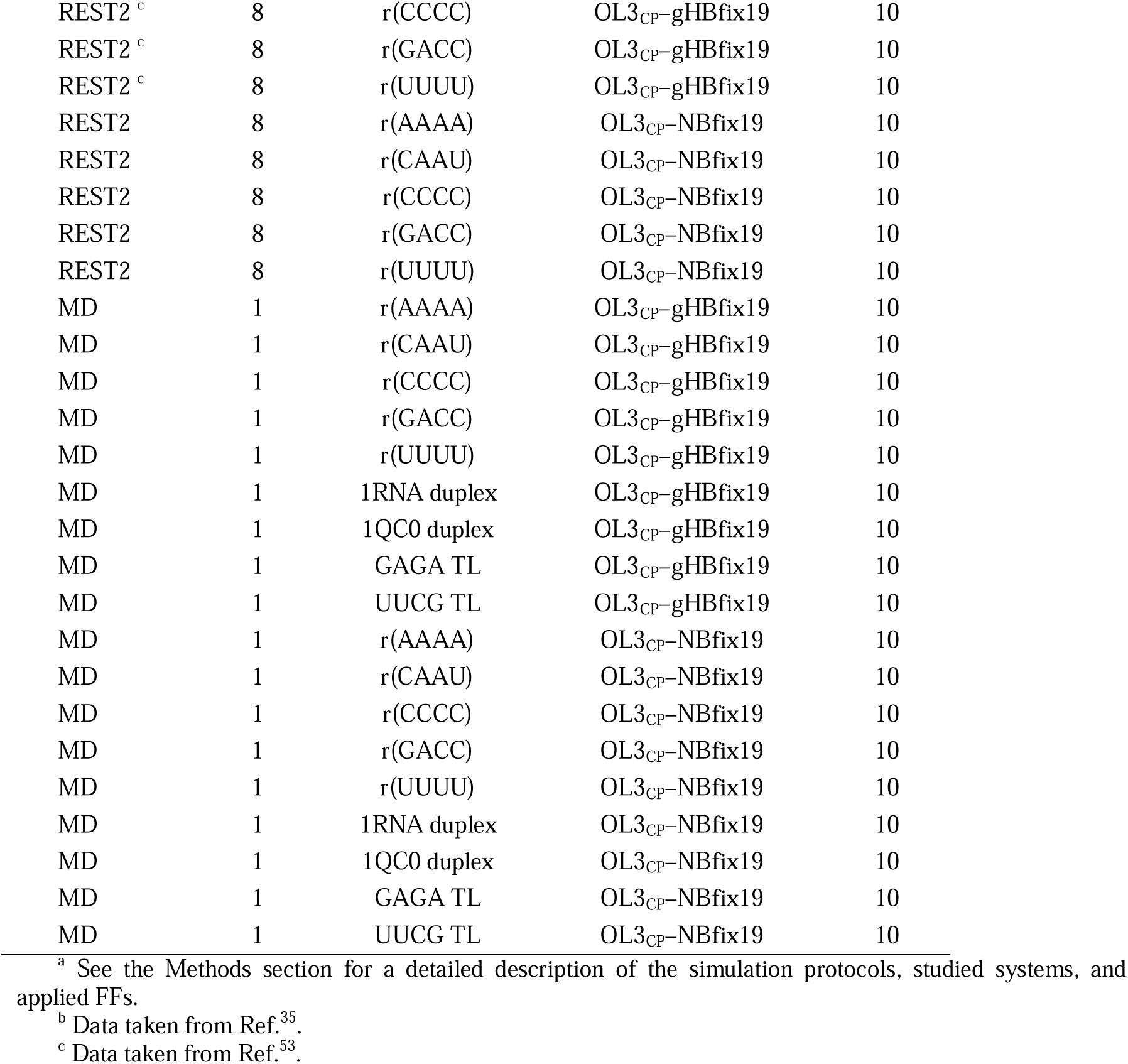
Overview of all enhanced sampling and standard MD simulations performed in this study.^a^.

The T-REMD training ensemble was generated using the OL3_CP_–gHBfix19 FF. All testing simulations were performed using either reference OL3_CP_–gHBfix19 FF or a newly derived OL3_CP_–NBfix19 FF, where the gHBfix19 terms were replaced by the corresponding NBfix reparameterizations introduced in this work. To validate individual NBfix reparameterizations, additional validation T-REMD simulations were carried out using *hybrid* FF variants (i.e., OL3_CP_–gHBfix19–NBfix_NH…N_, OL3_CP_–gHBfix19–NBfix_OH…nbO_, and OL3_CP_–gHBfix19–NBfix_OH…bO_), in which only a single interaction type originally modified by gHBfix19 was replaced by its NBfix counterpart, while all remaining gHBfix19 terms were retained. All systems underwent a standard equilibration protocol consisting of initial solvent minimization with positional restraints on RNA heavy atoms, followed by constant-pressure equilibration to stabilize the system density. The RNA solute was then minimized in several stages with gradually decreasing restraints on the sugar–phosphate backbone. Heating to 298 K was performed in two phases, first under constant-volume and subsequently under constant-pressure conditions.

### General MD settings

Systems were solvated in explicit OPC water^45^ using rectangular simulation boxes with a minimum solute–box distance of 10 Å. Monovalent ions were added to neutralize the systems and to achieve an ∼0.15 M KCl excess salt concentration. K^+^ and Cl^−^ ion parameters were taken from the Joung and Cheatham (KL: *R* = 1.590 Å, ε = 0.2795 kcal/mol; ClL: *R* = 2.760 Å, ε = 0.0117 kcal/mol).^46^ Long-range electrostatic interactions were treated with the particle mesh Ewald method^47, 48^ using a real-space cutoff of 10 Å and a grid spacing of 1 Å. Hydrogen mass repartitioning^49^ was applied to all simulations, enabling a 4 fs integration timestep. All standard MD simulations were performed for 10 μs (Table 1).

### T-REMD settings

All T-REMD^31^ simulations employed 64 replicas distributed over a temperature range of ∼278–461 K, selected to yield an average exchange probability of approximately ∼25 %. Simulations were carried out in the NVT ensemble under periodic boundary conditions.

Temperature was regulated using Langevin dynamics with a friction coefficient of 2 psL¹, and exchange attempts were carried out every 10 ps. Validation T-REMD simulations were run for 5 μs per replica when assessing individual NBfix substitutions, whereas the fully optimized OL3_CP_–NBfix19 FF was simulated for 10 μs per replica.

### REST2 settings

We also performed replica exchange solute tempering (REST2)^50^ simulations, which were performed at ∼298 K (the reference replica) with 8 replicas for all five TNs (Table 1). The scaling factor (λ) values ranged from 1 to 0.601700871, corresponding to the effective solute temperature range from ∼298 K to ∼500 K. Those values were chosen to maintain an exchange rate above 20%. REST2 simulations were performed with the AMBER18^41^ GPU MD simulation engine (pmemd.cuda).^51^ Further details about REST2 settings can be found elsewhere.^11^

### Conformational analysis

Dominant conformations of five TNs sampled during REST2 simulations were identified using the density-based clustering algorithm of Rodriguez and Laio^52^ combined with εRMSD.^39^ Because closely related conformational states with frequent interconversion are not be fully resolved by clustering alone, we complemented this analysis with a targeted εRMSD-based state assignment. Representative PDB structures of key conformations were used as references, and state populations were estimated from the trajectories using conservative εRMSD cutoffs. Detailed methodological descriptions are provided in our previous studies.^11, 17, 53^

### Comparison between MD and NMR data

The conformational ensembles of five TNs obtained from REST2 simulations were compared with previously published NMR experiments.^54–56^ We analyzed separately four NMR observables, i.e., (i) backbone ^3^J scalar couplings, (ii) sugar ^3^J scalar couplings, (iii) nuclear Overhauser effect intensities (NOEs), and (iv) the absence of specific peaks in NOE spectroscopy (uNOEs). Scalar couplings were calculated from instantaneous dihedral angles using standard Karplus relationships. NOE and uNOE intensities were obtained by ensemble averaging over the reference (unbiased) trajectories from REST2 simulations, applying the conventional inverse sixth-power distance dependence and subsequent back-transformation to effective distances.^32, 53^

For the observed quantities (scalar couplings and NOEs), agreement between simulation and experiment was quantified using a χ² metric. As in previous studies,^15, 32, 53^ the agreement with uNOEs, which represent the absence of experimentally detectable NOE peaks and therefore correspond to inequality constraints rather than direct measurements, was evaluated using a one-sided penalty function that quantifies violations of the corresponding distance limits. Because this quantity does not follow a χ² distribution, we report it separately in this study as a uNOE violation score *p*. In contrast to earlier studies where individual contributions were combined by averaging over all NMR signals including the unobserved NOEs,^15, 32, 53^ we here report the overall agreement between simulation and experiment *S*_total_ as the arithmetic mean of the four observable classes (backbone ³J couplings, sugar ³J couplings, NOEs, and uNOE constraints). This procedure avoids an artificial statistical bias arising from the typically larger number of uNOE constraints and ensures comparable contributions of the different experimental observables to the final agreement metric. Lower values of the individual χ² and *p* scores and of the overall metric *S*_total_ indicate better agreement between the simulated ensembles and experimental measurements.

## RESULTS&DISCUSSION

The goal of this study was to develop an NBfix-based parametrization that reproduces the effects of the gHBfix19 potential within the OL3_CP_–gHBfix19 RNA FF. The gHBfix formalism provides a flexible and physically transparent way to fine-tune specific non-bonded H-bond interactions, but its implementation is computationally demanding and requires additional restraint definitions for each donor–acceptor pair. To enable routine use in large-scale simulations, we sought to develop a methodology to transfer the physical effects of the gHBfix potential into a functionally equivalent set of NBfix terms that modify the LJ parameters for selected atom-type pairs outside the standard Lorentz–Berthelot combination rules. This approach preserves the underlying physics of the reference FF, in this particular case the OL3_CP_–gHBfix19 RNA FF, while achieving the same thermodynamic balance through a simpler and fully native non-bonded formulation compatible with standard AMBER FFs.

### Reweighting-Driven Reparameterization of gHBfix19 into NBfix Terms

To identify NBfix parameters reproducing the effect of gHBfix19 corrections, we employed a reweighting approach based on an equilibrated ensemble of the GAGA TL obtained from T-REMD simulation. As the training set, we used the final 5 μs of the 25 μs T-REMD trajectory initiated from the folded state, which was previously shown to be fully converged with respect to folding–unfolding equilibrium.^35^ This ensemble provided the relevant set of donor–acceptor distances corresponding to all H-bonds affected by the gHBfix19 terms.^11^ For every tested NBfix parameter combination, we recalculated the non-bonded energies and reweighted the ensemble to estimate the resulting folded-state population and the reliability of the reweighting, expressed by the *P*_eff_ values (see Methods). This procedure allowed us to systematically explore the LJ parameter space (radius (*R*) and well depth (ε)) for each interaction type and to select optimal values that best reproduce the original thermodynamic balance provided by the reference OL3_CP_–gHBfix19 FF while maintaining physical consistency with the parent OL3 FF.

Because the *P*_eff_ values of reweighting decreases with increasing deviation from the reference FF, we reparameterized the gHBfix19 terms individually. The gHBfix19 potential was introduced to correct two key deficiencies of the OL3 FF: (i) the underestimated stability of base–base H-bonds, addressed by adding +1.0 kcal/mol stabilization to all –NH···N– interactions, and (ii) the overstabilized ribose–phosphate contacts, reduced by penalizing –OH···(n)bO– interactions by −0.5 kcal/mol.^11^ Since the –NH···N– term plays a central role in stabilizing canonical and noncanonical base pairing, we addressed it first. Although gHBfix19 is formally applied to the – H···N– donor–acceptor pairs, the corresponding interaction can in principle be captured either through an NBfix modification of the heavy-atom LJ interaction (–N···N–) or of the H–acceptor interaction (–H···N–). To identify which representation best reproduces the original gHBfix19 behavior, we performed two-dimensional reweighting scans over the LJ radius *R* and well depth ε for both –N···N–and –H···N– pairs using the equilibrated T-REMD ensemble of the GAGA TL.

Reweighting demonstrates that only modifications of the –H···N– LJ pair can simultaneously recover the correct folded-state population and maintain acceptable reliability (Figure 1). In contrast, tuning the heavy-atom –N···N– interaction fails to recover the correct thermodynamic balance and yields negligible *P*_eff_ values of the reweighting (Figure S1 in the Supporting Information). Optimal parameters were selected to reproduce the folded fraction obtained with the reference OL3_CP_–gHBfix19 FF, while maximizing the reliability. The best-performing parameters were found at *R* = 2.1 Å and ε = 0.63 kcal/mol, compared to the default OL3 values of *R* = 2.424 Å and ε = 0.0517 kcal/mol. This represents a modest strengthening and slight contraction from the original OL3 FF values, i.e., a larger ε and slightly smaller *R*, consistent with the intended +1.0 kcal/mol stabilization introduced by gHBfix19 potential. These parameters were applied to the LJ pair between H atoms of amino or imino groups (AMBER atom type H) and N acceptors capable of forming H-bonds (AMBER atom types NB and NC, corresponding to purine N7 and pyrimidine N3/N1 atoms, respectively). The comparison of the standard OL3 FF, reference OL3_CP_–gHBfix19 FF, and the newly optimized potential (Figure 1) illustrates that the new NBfix parametrization successfully reproduces the intended stabilization of –NH···N– interactions without altering the overall shape of the non-bonded potential. A slight shift in the position of the LJ minimum is nevertheless observed for the fitted OL3_CP_–NBfix19 FF relative to the original OL3_CP_–gHBfix19 profile (Figure 1C). To assess potential site-specific effects of this shift, we calculated radial distribution functions *g*(r) for –NH···N–interactions in canonical RNA duplexes. The resulting profiles are nearly identical, indicating no detectable differences between the two FFs (Figure S2 in the Supporting Information).

**Figure 1.**
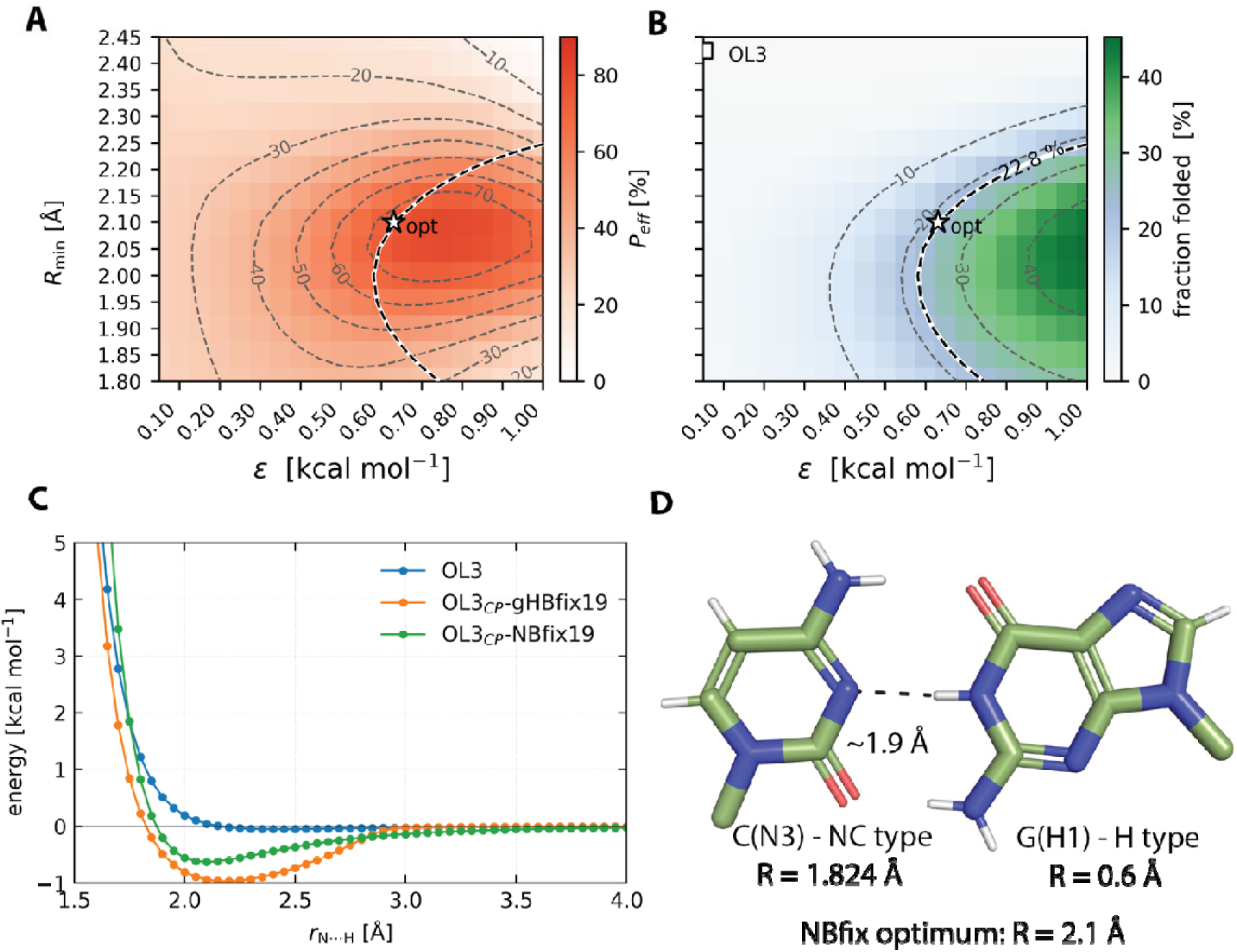
Reweighting-based optimization of NBfix parameters for the –H···N– interaction. (A) Normalized effective sample size *P*_eff_ of the reweighted ensemble as a function of LJ radius *R* and well depth *ε*. (B) Reweighted folded-state fraction over the same range of parameters, with the dashed contour denoting the target folded-state population from the converged OL3_CP_–gHBfix19 T-REMD simulation. The star indicates the optimal NBfix parameters, and the square the original OL3 values. (C) LJ interaction profiles for the –H···N–pair in OL3, OL3_CP_–gHBfix19, and the optimized NBfix model. (D) Canonical GC base pair showing the AMBER van der Waals radii for the C(N3) and G(H1) atoms, their typical H-bond distance, and the optimized NBfix radius.

In analogy to the –NH···N– reparameterization, we next examined the reparametrization of the – OH···nbO– and –OH···bO– gHBfix19 terms that penalized each of these interactions by −0.5 kcal/mol. The OL3 (AMBER) FF assigns zero LJ parameters to H atoms of 2’-OH groups, meaning that the repulsive part of the potential is entirely missing and the FF relies on the repulsion between the corresponding heavy atoms. This deficiency is reflected in previous observations of overstabilized ribose–phosphate interactions^10, 11, 21, 22, 57^ and is further highlighted by the reweighted maps, which show that eliminating the repulsive term (ε → 0) leads to poor *P*_eff_ values and incorrect thermodynamic balance (Figure 2). We therefore focused on NBfix parametrizations that introduce a physically meaningful repulsive component to the –H···O– interaction while reproducing the folded-state population of the reference OL3_CP_–gHBfix19 FF and preserving sufficient *P*_eff_ values. Because unconstrained optimization tended to drive ε toward zero compensated by *R* reaching unrealistically large values, we fixed the well depth at a small but finite value of ε = 0.01 kcal/mol and optimized only the LJ parameter *R*. The resulting optimal parameters are *R* = 2.73 Å for the –H···nbO–interaction and *R* = 3.03 Å for the –H···bO– interaction. These values reintroduce the missing steric repulsion between H-atoms from 2’-OH groups and phosphate oxygens while accurately capturing the weakening effect originally imposed by the gHBfix19 potential.^11^

**Figure 2.**
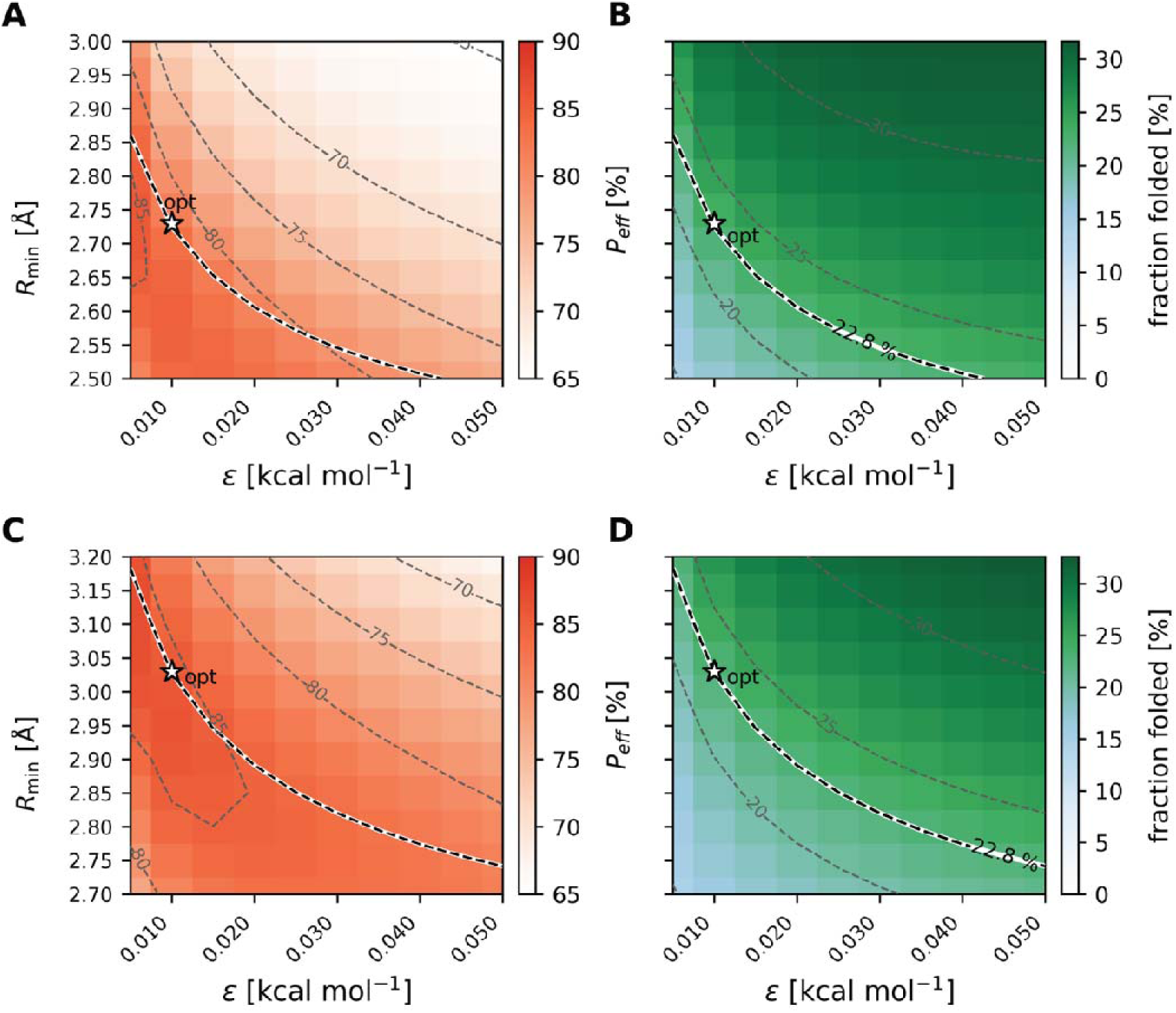
Reweighting-based optimization of NBfix parameters for the –H···(n)bO– interactions. Pannels (A) and (C) show *P*_eff_ values of the reweighted ensembles as a function of *R* and *ε* for the –H···nbO– and –H···bO–interactions, respectively. Pannels (B) and (D) correspond to reweighted folded-state fractions, with dashed contours marking the target value obtained from the converged OL3_CP_–gHBfix19 T-REMD simulation. Stars indicate optimal NBfix parameters; the original OL3 FF values (no modification) fall outside the plotted range.

### Validation of gHBfix19 Reparameterization via NBfix

Reweighting provides an efficient way to identify NBfix parameters that should reproduce the energetic effects of the gHBfix19 corrections, and high *P*_eff_ values indicate that the reweighted ensemble remains statistically representative of the reference simulation. Nevertheless, any such parametrization must be validated by explicit simulations. Because the three gHBfix19 terms (–NH···N–, –OH···nbO–, and –OH···bO–) were optimized individually, we performed three corresponding validation simulations in which each interaction type was selectively removed from the gHBfix19 and replaced by its optimized NBfix counterpart. For each case, we initiated a 64-replica T-REMD simulation of 5 μs per replica using the final ensemble from the original 25 μs-long OL3_CP_–gHBfix19 T-REMD run as starting structures. In all three tests, the folded-state population remained close to the reference value of 22.8 % obtained with OL3_CP_–gHBfix19 FF. Replacing the –NH···N– term yielded a folded fraction of 30.4 % ± 1.2 % (mean and standard deviation of fraction folded over final 2 μs derived from the *ε*RMSD-based definition with cutoff of 0.7), while substitutions of the –OH···nbO– and –OH···bO– terms gave 26.4 % ± 5.5 % and 17.4 % ± 3.8 % folded populations, respectively (Figure 3; see Figures S3-S5 in Supporting Information for time evolution of main structural states). Thus, two parametrizations slightly increased the folded-state population relative to the reference, whereas the –OH···bO–substitution led to a modest decrease, all remaining within acceptable deviations for a partial term-by-term replacement.

**Figure 3.**
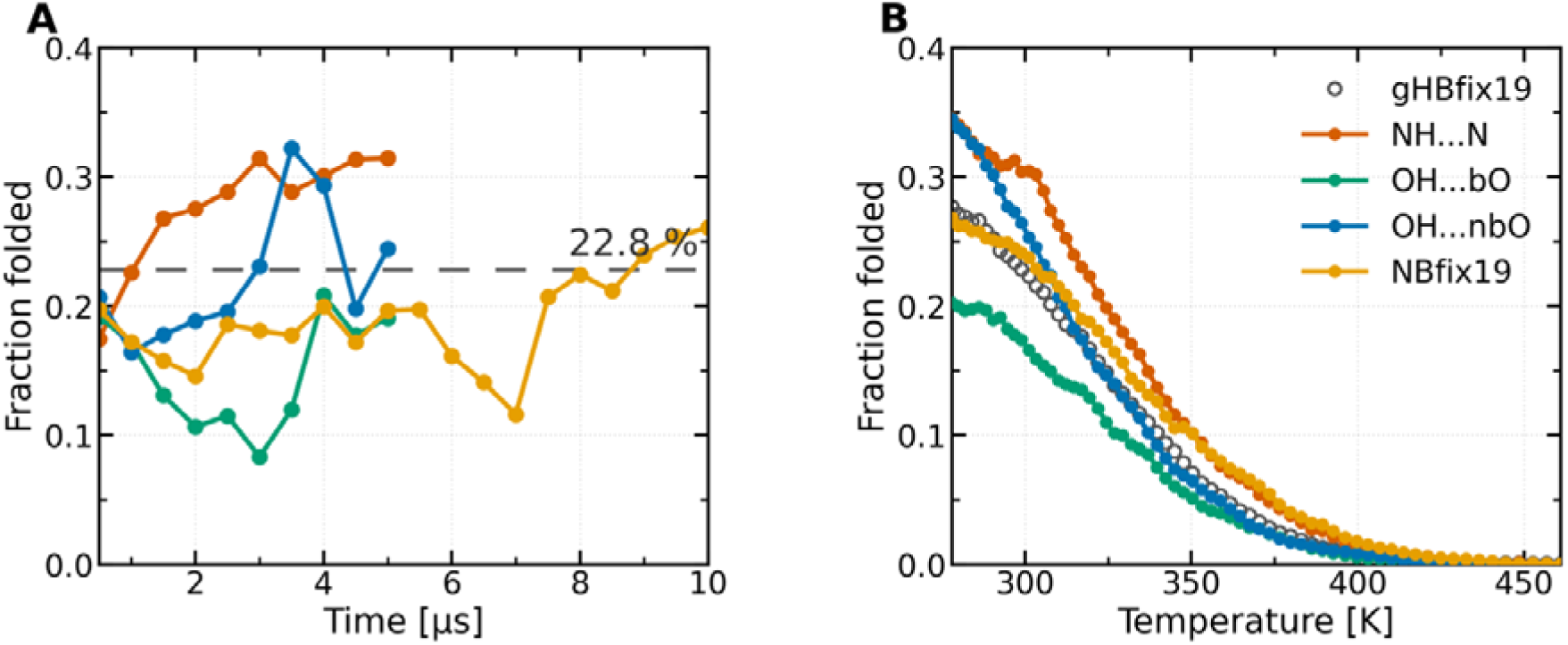
Validation of NBfix reparameterizations through 64-replica T-REMD simulations, either substituting the individual gHBfix19 terms with their corresponding NBfix counterparts or using the full NBfix model replacing all gHBfix19 corrections, i.e., the OL3_CP_–NBfix19 FF. (A) Time evolution of the folded-state population at 298.996 K. (B) Folded-state fractions averaged over the final 2 μs of each simulation including the OL3_CP_–gHBfix19 reference as a function of temperature.

As a final test of reweighting-driven NBfix parameter optimization, we carried out a T-REMD simulation in which all three gHBfix19 components were simultaneously removed and replaced by their optimized NBfix parametrizations, yielding a fully gHBfix-free variant (the OL3_CP_–NBfix19 FF). This test evaluates not only the accuracy of individual NBfix substitutions but also their collective behavior and potential cross-interactions that cannot be detected in isolated reweighting scans or single-term validation runs. Starting from the same equilibrated structures used in the partial tests, the simulation was propagated for 10 μs. Over the final 2 μs, the folded-state population stabilized at 23.8% ± 2.0 %, which is in close agreement with the reference folded fraction of 22.8 % obtained with the original OL3_CP_–gHBfix19 FF (Figure 3; see Figure S6 in Supporting Information for time evolution of main structural states). This confirms that the jointly applied NBfix terms do not introduce adverse cooperative effects and that the complete NBfix reformulation accurately preserves the intended balance between folded and unfolded ensembles.

### OL3_CP_–NBfix19 FF Benchmarking Across Representative RNA Motifs

To evaluate the transferability of the newly derived OL3_CP_–NBfix19 FF beyond the reference training system, we assessed its performance across three representative classes of RNA motifs of increasing structural complexity: (i) short TNs, (ii) canonical A-form duplexes, and (iii) TL hairpins (see Methods and Table 1). The results obtained with the new NBfix-based FF were systematically compared with those obtained using the reference OL3_CP_–gHBfix19 FF.

We first examined a set of five RNA TNs (r(AAAA), r(CAAU), r(CCCC), r(GACC), and r(UUUU)), which represent highly flexible benchmark systems with NMR reference data. Standard MD simulations (10 μs-long) were initially performed for each TN using both OL3_CP_–gHBfix19 and OL3_CP_–NBfix19 FFs. Despite the relatively long simulation times for such small systems, these trajectories did not yield fully converged conformational ensembles. The simulations remained dominated by the initial A-form–like conformations together with partially stacked and intercalated states (Tables S1 and S2 in Supporting Information). Transitions between these states occurred on the hundreds-of-ns to μs timescale, and their relative populations varied substantially between independent trajectories (Figures S7-S11 in Supporting Information). These observations highlight the intrinsically slow conformational exchange of TN systems and indicate that even multi-μs conventional MD simulations are insufficient to reliably sample their equilibrium ensembles.

To achieve better conformational sampling, we therefore employed REST2 enhanced sampling simulations for all five TNs. REST2 substantially improved transitions among major conformational states and enabled a more reliable redistribution of populations. Both OL3_CP_–NBfix19 and OL3_CP_–gHBfix19 FFs produced qualitatively similar ensembles with comparable populations of dominant conformers (Table 2 and Table S3 in Supporting Information), including the native A-major states and the well-known spurious intercalated structures.^10, 11, 18, 32, 53, 56–58^ Importantly, both FFs also yielded similar levels of agreement with the available NMR observables, as reflected by comparable χ² and *p* values for the individual observable classes and by the overall *S*_total_ metric (Table 2). Overall, the OL3_CP_–NBfix19 FF reproduces the conformational behavior of the TN set at a level comparable to the reference OL3_CP_–gHBfix19 FF.

**Table 2:** Overview of REST2 simulations of RNA TNs.^a^.

| TNs | Observable |  | OL3<sub>CP</sub>-gHBfix19<sup>b</sup> | OL3<sub>CP</sub>-NBfix19 |
| --- | --- | --- | --- | --- |
| r(AAAA) | Clusters [%]<sup>c</sup> | A-form | ~29 | ~29 |
|  |  | Intercalated | ~11 | ~9 |
| | NMR comparison ($\chi^2$)<sup>d</sup> | <sup>3</sup>J-backbone | 0.66 | 0.66 |
|  |  | <sup>3</sup>J-sugar | 1.23 | 1.05 |
|  |  | NOE | 1.08 | 1.20 |
|  |  | uNOE | 1.14 | 1.07 |
| | | $S_{\text{total}}$ | 1.03 | 1.00 |
| r(CAAU) | Clusters [%]<sup>c</sup> | A-form | ~33 | ~50 |
|  |  | Intercalated | ~51 | ~31 |
| | NMR comparison ($\chi^2$)<sup>d</sup> | <sup>3</sup>J-backbone | 0.73 | 0.58 |
|  |  | <sup>3</sup>J-sugar | 1.14 | 0.92 |
|  |  | NOE | 1.81 | 1.34 |
|  |  | uNOE | 6.07 | 4.44 |
| | | $S_{\text{total}}$ | 2.44 | 1.82 |
| r(CCCC) | Clusters [%]<sup>c</sup> | A-form | ~55 | ~71 |
|  |  | Intercalated | ~40 | ~23 |
| | NMR comparison ($\chi^2$)<sup>d</sup> | <sup>3</sup>J-backbone | 0.44 | 0.35 |
|  |  | <sup>3</sup>J-sugar | 0.68 | 0.47 |
|  |  | NOE | 1.54 | 1.27 |
|  |  | uNOE | 3.22 | 1.98 |
| | | $S_{\text{total}}$ | 1.47 | 1.02 |
| r(GACC) | Clusters [%]<sup>c</sup> | A-form | ~74 | ~87 |
|  |  | Intercalated | ~2 | ~0 |
| | NMR comparison ($\chi^2$)<sup>d</sup> | <sup>3</sup>J-backbone | 0.39 | 0.34 |
|  |  | <sup>3</sup>J-sugar | 0.50 | 0.44 |
|  |  | NOE | 1.13 | 1.20 |
|  |  | uNOE | 0.34 | 0.01 |
| | | $S_{\text{total}}$ | 0.59 | 0.50 |
| r(UUUU)<sup>e</sup> | Clusters [%]<sup>c</sup> | Unstructured | ~3 | ~16 |
|  |  | 1-3/2-4 stack | ~38 | ~31 |
| | NMR comparison ($\chi^2$)<sup>d</sup> | <sup>3</sup>J-backbone | 0.53 | 0.50 |
|  |  | <sup>3</sup>J-sugar | 0.87 | 0.80 |
|  |  | NOE | 3.53 | 4.40 |
|  |  | uNOE | 1.65 | 2.08 |
| | | $S_{\text{total}}$ | 1.65 | 1.95 |

<sup>a</sup> All simulations used 8 replicas and were run for 10 μs (the last 7 μs were used for data analysis). See the Methods section for a detailed description of the REST2 protocol, studied TNs, and applied FFs.
<sup>b</sup> Data taken from the Ref.<sup>53</sup>.
<sup>c</sup> Only main clusters (typically native A-form and spurious intercalated states) are listed. See Supporting Information for complete results from the clustering analysis.
<sup>d</sup> The agreement between simulation and experiment was quantified using $\chi^2$ values comparing calculated and experimental backbone <sup>3</sup>J scalar couplings, sugar <sup>3</sup>J scalar couplings, and NOEs, and using the uNOE violation score $p$ . The overall agreement metric $S_{\text{total}}$ was calculated as the arithmetic mean of the four observable classes. See Methods for details.
<sup>e</sup> The UUUU TN is expected to populate diverse conformations rather than a canonical A-RNA structure;<sup>56</sup> therefore, we report populations of unstructured states and 1-3/2-4 stacking arrangements.

Next, we evaluated the stability of canonical A-form RNA helices using two duplex systems: the r(GCACCGUUGG)L decamer derived from the high-resolution crystal structure 1QC0^42^ and the alternating r(UUAUAUAUAUAUAA)L duplex derived from the 1RNA^43^ structure. In contrast to the TN systems discussed above, which required enhanced sampling to converge their conformational ensembles, the duplex simulations were designed primarily to assess the stability and local dynamics of the native A-form structure. In both cases, 10 μs standard MD simulations with the OL3_CP_–NBfix19 FF maintained stable A-form helices without detectable structural distortions. Global helical parameters and local base-pair geometries remained within the expected ranges for canonical A-RNA and were consistent with those obtained using the reference OL3_CP_–gHBfix19 FF (Tables S4 and S5 in Supporting Information). Terminal base-pair opening (fraying) occurred occasionally at duplex ends but showed no systematic differences between the two tested FF variants (Table S5 in Supporting Information). These results indicate that the NBfix-based FF preserves the stabilizing effects of the gHBfix19 correction, which was previously shown to substantially reduce excessive base-pair fraying observed in earlier OL3 simulations lacking this correction.^11^

Finally, we assessed the behavior of two TL systems representing compact RNA motifs with well-defined tertiary interactions: the GAGA TL used as the reference training system and the UUCG TL as an independent validation case (see Methods). In 10 μs standard MD simulations, the GAGA TL remained stably folded in both OL3_CP_–NBfix19 and OL3_CP_–gHBfix19 FFs, exhibiting low RMSD values and persistent sampling of the native loop conformation (Figures S12 and S13 in Supporting Information). This result is consistent with the T-REMD validation described above and confirms that the newly derived OL3_CP_–NBfix19 FF preserves the thermodynamic balance of this reference system. In contrast, the UUCG TL lost its native loop conformation in both simulations. Disruption of the signature loop interactions occurred within the μs timescale in both simulations, leading to loss of the native loop conformation (Figures S14 and S15 in Supporting Information). The A-form stem containing five canonical base pairs remained stable in both simulations for entire 10 μs-long timescale. The limited stability of the UUCG TL is consistent with previous studies showing that this motif remains challenging for the majority of current RNA FFs, including more recent parametrizations and polarizable models.^11, 16, 23^ We also tested the latest machine-learning–assisted refinement of the OL3 FF incorporating experimental data, i.e., the OL3-vdW7-PAK version.^27^ Rather surprisingly, it showed comparably limited performance for the UUCG TL (see Supporting Information for full details). Notably, the original gHBfix19 correction itself was not expected to fully stabilize this motif; previous work indicated that only the later gHBfix21 version^19^ approaches near-zero folding free energies for the 8-mer UUCG TL, whereas most currently available RNA FFs still predict substantially positive folding free energies for the native state.^16^

Taken together, these benchmark tests demonstrate that the OL3_CP_–NBfix19 FF preserves the key structural and thermodynamic characteristics of the reference OL3_CP_–gHBfix19 parametrization across diverse RNA motifs, while providing a simplified and computationally efficient representation of the underlying H-bond corrections.

## CONCLUDING REMARKS

In this work, we introduced a systematic strategy for translating externally defined H-bond correction potentials into native FF terms. Using a quantitatively converged T-REMD ensemble of the GAGA TL as a reference system, we employed a statistically controlled reweighting protocol to identify NBfix LJ parameters that reproduce the thermodynamic effects of the gHBfix19 corrections^11^ within the OL3_CP_ RNA FF. Reweighting approaches have previously been employed to derive correction terms for MD FFs (see, e.g., Refs.^19, 59–62^). In the present case, the task was facilitated by the availability of a previously optimized set of corrections, that were obtained by trial-and-error optimization,^11^ which we reimplemented here using a different functional form. This allowed us to use the pre-optimized ensemble as a reference, thereby alleviating issues associated with insufficient statistical overlap in reweighting, which would otherwise require computationally demanding iterative procedures.^63^ At the same time, the use of a standard LJ parametrization presents specific challenges. While LJ potentials are efficiently implemented in most MD codes, making the resulting parametrization readily transferable, their steep repulsive region at short distances reduces the statistical efficiency of the reweighting procedure compared to smoother H-bond correction potentials.^19^ Because the statistical reliability of reweighting decreases with increasing deviation from the reference Hamiltonian, the reparameterization had to be carried out sequentially, addressing individual gHBfix19 components step by step. This approach is conceptually related to that proposed in Ref.^60^, where reweighting was performed for individual amino acids and the combined effect of the corrections was evaluated *a posteriori*. The resulting fully reformulated variant, denoted OL3_CP_–NBfix19, reproduces the folding equilibrium of the reference system and exhibits performance comparable to the original OL3_CP_–gHBfix19 FF across a representative set of RNA motifs. Importantly, this variant can be readily implemented in standard MD packages such as AMBER^41^ and GROMACS,^64^ without the need for additional computationally expensive correction terms as required in Ref.^19^.

More broadly, the present work highlights a practical strategy for addressing a long-standing challenge in FF development. Non-bonded interactions remain one of the major sources of residual imbalance in contemporary RNA FFs,^10, 11, 17, 19, 21, 22, 25, 27, 57, 61, 65–69^ yet their direct reparameterization is notoriously difficult due to strong coupling between van der Waals parameters, electrostatics, and other FF terms. Targeted correction potentials such as gHBfix provide a powerful and physically transparent way to tune specific interaction classes without introducing widespread side effects, making them highly effective tools for diagnosing and refining problematic interactions. However, their reliance on additional external potentials and explicit donor–acceptor restraints complicate their routine use in large-scale production simulations. By demonstrating how the effects of such targeted corrections can be quantitatively transferred into native NBfix parameters, our approach bridges the gap between efficient FF tuning and practical production-level FF implementations.

Finally, the workflow presented here establishes a general framework for reweighting-driven refinement of non-bonded FF parameters. While the present study focused on the relatively compact gHBfix19 correction scheme, the same strategy can in principle be applied to more complex correction frameworks or larger sets of interaction classes such as gHBfix21.^19^ In such cases, automated optimization procedures and machine-learning–assisted parameter exploration will likely become necessary to efficiently explore the high-dimensional parameter space. Nevertheless, the present results demonstrate that targeted interaction corrections can be systematically translated into efficient native FF parametrizations while preserving the intended thermodynamic balance of RNA conformational ensembles.

## Supporting information

Supporting Information for the article

## ASSOCIATED CONTENT

### Supporting Information

The Supporting Information is available free of charge via the Internet at http://pubs.acs.org/ and contains parmed input file for the NBfix19 modification of the OL3_CP_ FF, details about additional simulations with the OL3-vdW7-PAK FF, supporting tables and figures.

### Data Availability Statement

All raw MD data are available on Lexis platform: https://portal.lexis.tech (project exa4mind_wp4).

## ACKNOWLEDGMENT

This work was supported by the Czech Science Foundation (grant number 23-05639S; to V.M., and J.Š.). This research also received the support of EXA4MIND, a European Uniońs Horizon Europe Research and Innovation programme under grant agreement N° 101092944 (V.M., M.O. and P.B). Views and opinions expressed are however those of the author(s) only and do not necessarily reflect those of the European Union or the European Commission. Neither the European Union nor the granting authority can be held responsible for them. P.K., M.O. and P.B. were supported by ERDF/ESF project TECHSCALE (No. CZ.02.01.01/00/22_008/0004587). G.B. acknowledges the Italian National Centre for HPC, Big Data, and Quantum Computing (grant No. CN00000013), founded within the Next Generation EU initiative. This work was also supported by the Ministry of Education, Youth and Sports of the Czech Republic through the e-INFRA CZ (ID:90254).

