## Supporting Information for the article for "From gHBfix to NBfix: Reweighting-Driven Refinement of Hydrogen-Bond Interactions in RNA Force Fields"

### Table of Contents

SUPPORTING RESULTS ........ - 2 -

SUPPORTING TABLES ........ - 4 -

SUPPORTING FIGURES ........ - 7 -

REFERENCES ........ - 16 -

SUPPORTING RESULTS

**ParmEd file implementing the NBfix19 modification for the OL3$_{CP}$ RNA Force Field.**[1-6]

ParmEd File Usage: $AMBERHOME/bin/parmed topologyfile_OL3CP.parm7 -i parmed_NBfix19.in

parmed_NBfix19.in

```
#
# parmed input file to modify off-diagonal Lennard-Jones parameters in order to replace external gHBfix19 H-bond correction potential (Kuehrova & Mlynsky et al., JCTC 2019) with the NBfix19 modification
# for details, see Mlynsky V., Kuehrova P., Bussi G., Otyepka M., Sponer J., and Banas P., 2026
#
# IMPORTANT: this assumes that the topology was built with OL3 RNA force field (Zgarbova et al., JCTC 2011) further adjusted by the phosphate oxygen van der Waals correction (CP) by Steinbrecher et al., JCTC 2012
# other force fields may have different atom types and need different atom masks below
#
#
# NBfix modification of -NH...N- interactions according to gHBfix19 potential
# all A_N6(H61/H62), C_N4(H41/H42), G_N2(H21/H22), G_N1(H1), and U_N3(H3) hydrogens share same and unique atom type H
# A(N1), A(N3), A(N7), C(N3), G(N3), and G(N7) share same NB and NC atom types, but these are also share with, e.g., N, NA, N2 .., so we need to split it
addLJType @%NB,NC
changeLJPair @%NB,NC @%H 2.1 0.63
#
# NBfix modification of -OH...bO/nbO- interactions according to gHBfix19 potential
# all 2'-OH hydrogens (i.e., HO2', HO3', and HO5') share same and unique atom type HO
# all bridging (bO; O5' and O3') and nonbridging (nbO; O1P and O2P) phosphate oxygens share same and unique atom types OR and OP
changeLJPair @%HO @%OP 2.73 0.01
changeLJPair @%HO @%OR 3.03 0.01
#
#
# CHANGE the newtopologyfile to your new name to output a parm7 file with the NBfix19 modification
outparm newtopologyfile_NBfix19.parm7
```

**Additional UUCG tetraloop simulations with newly derived reparameterization of the OL3 Force Field.** During completing results for this work, machine-learning–assisted reparameterization of the OL3 Force Field (FF), which incorporates experimental data into fitting schemes, has been published by Pak and coworkers (denoted here as OL3-vdW7-PAK)[7]. The authors presented very encouraging results considering 10-mer r(cacUUCGgug) tetraloop (UUCG TL) derived from the 14-mer NMR structure (PDB ID 2KOC[8]), showing dominant population of the native state during multiple-walker on-the-fly probability (OPES)[9, 10] enhanced-sampling simulations.[7]

Here, we performed a set of four independent 10 μs-long standard MD simulations for both shorter r(gcUUCGgc) 8-mer and longer r(cacUUCGgug) 10-mer TLs with the OL3-vdW7-PAK FF. Starting structures were taken from our previous work[11] and excised from the NMR structure (snapshot #13)[8] for the shorter 8-mer and longer 10-mer TL, respectively. Simulations were performed in Gromacs2020[12] with FF libraries published by the original authors (directly downloaded from the Zenodo repository: https://zenodo.org/records/15622306).[7] Systems were solvated with the OPC3 water model.[13] KCl ions described by Li and Merz parameters[14] for the OPC3 water were added to neutralize the system and to establish excess-salt ion concentration of 0.15 M. Gromacs simulations were performed in a rhombic dodecahedral box and bonds involving hydrogens were constrained using the LINCS algorithm.[15] The cut-off distance for the direct space summation of the electrostatic interactions was 10 Å and the simulations were performed at 298 K using the stochastic velocity rescale thermostat.[16]

Surprisingly, our results from the standard MD simulations showed that the UUCG TL is very poorly described by the OL3-vdW7-PAK FF, in sharp contrast with the original paper.[7] We observed almost immediate and irreversible loss of the signature interactions with the loop within first few hundreds of ns in all four attempted MD simulations of the shorter 8-mer TL (Figure S16). Comparable behavior was observed during simulations of the longer 10-mer TL, where the stability of the native state was slightly prolonged towards ~2 μs (in 3 out of 4 MD simulations; Figure S17). The A-form stem containing two (8-mer) or three (10-mer) canonical base pairs remained stable in all simulations for the entire 10 μs-long timescale. The irreversible disruption of the native state was initiated by the key G nucleotide (G6 and G7 for 8-mer and 10-mer, respectively) of the loop with *syn* conformation of $\chi$ dihedral, which left the binding site and flipped into *anti* state. This is typical and well documented disruption pathway of contemporary RNA FFs.[17, 18]

### SUPPORTING TABLES

**Table S1.** Clustering analysis of 10 μs-long standard MD simulations of five TNs.[a]

| System | Cluster | $OL3_{CP}$–gHBfix19 | $OL3_{CP}$–NBfix19 |
|---|---|---|---|
| r(AAAA) | A-form | 28.9 | 25.6 |
| | Intercalated | - | 1.0 |
| | Bulge-out | 15.5 | 2.2 |
| | Flipped | 24.3 | 34.9 |
| | 1-2/3-4 stack | 1.3 | - |
| | Loop | 1.5 | - |
| | 1-3/2-4 stack | - | 9.7 |
| r(CAAU) | A-form | 29.7 | 47.6 |
| | Intercalated | 24.5 | - |
| | Flipped | - | 3.9 |
| | 1-2/3-4 stack | - | 1.1 |
| | Bulge-out | 23.2 | 31.0 |
| r(CCCC) | A-form | 81.5 | 65.8 |
| | Intercalated | - | 24.5 |
| | Bulge-out | 6.9 | 2.0 |
| | 1-3/2-4 stack | - | 3.4 |
| r(GACC) | A-form | 88.0 | 80.5 |
| | Loop | 4.4 | - |
| | Bulge-out | - | 9.5 |
| r(UUUU) | A-major | 4.4 | 6.8 |
| | Intercalated | 4.3 | 3.4 |
| | 1-2/3-4 stack | 2.0 | 7.4 |
| | 1-3/2-4 stack | 34.0 | 29.8 |
| | Bulge-out | 23.4 | 19.4 |
| | Loop | 9.7 | 2.6 |
| | Flipped | 1.4 | - |

[a] See Methods in the main text for details about clustering algorithm and Ref.[19] for details about structural states.

**Table S2.** Complementary clustering analysis of 10 μs-long standard MD simulations of five TNs.[a]

| System | Cluster | $OL3_{CP}$–gHBfix19 | $OL3_{CP}$–NBfix19 |
|---|---|---|---|
| r(AAAA) | A-major | 12.5 | 15.2 |
| | A-minor | 3.8 | 4.2 |
| | Intercalated | 0.1 | 0.0 |
| r(CAAU) | A-major | 17.5 | 30.5 |
| | A-minor | 6.1 | 10.1 |
| | Intercalated | 22.3 | 0.7 |
| r(CCCC) | A-major | 58.0 | 54.9 |
| | A-minor | 8.2 | 5.0 |
| | Intercalated | 1.8 | 24.3 |
| r(GACC) | A-major | 67.4 | 47.3 |
| | A-minor | 8.1 | 6.1 |
| | Intercalated | 0.2 | 3.2 |
| | Stack | 0.0 | 6.1 |
| | Loop | 4.3 | 0.0 |
| r(UUUU) | A-major | 4.2 | 6.5 |
| | A-minor | 13.1 | 9.5 |
| | Intercalated | 3.4 | 2.4 |

[a] Targeted $\varepsilon$RMSD-based state assignment was applied for predefined cluster types, where representative PDB structures of key conformations were used as references (see Methods in the main text for details and Ref.[19] for details about structural states).

**Table S3.** Clustering analysis of REST2 simulations of five TNs.[a]

| System | Cluster | $OL3_{CP}$–gHBfix19 | $OL3_{CP}$–NBfix19 |
|---|---|---|---|
| r(AAAA) | A-form | 28.8 | 28.7 |
| | Intercalated | 11.3 | 8.9 |
| | Bulge-out | 8.8 | 3.7 |
| | Flipped | 20.5 | 39.2 |
| | 1-2/3-4 stack | 4.6 | - |
| | 1-3/2-4 stack | 3.9 | 2.2 |
| r(CCCC) | A-form | 55.3 | 71.1 |
| | Intercalated | 40.4 | 23.3 |
| | Bulge-out | - | 1.1 |
| r(UUUU) | A-form | - | 8.7 |
| | 1-3/2-4 stack | 31.4 | 38.2 |
| | Bulge-out | 12.1 | 18.3 |
| | Flipped | 13.3 | 12.7 |
| | Unstructured | 15.5 | 3.3 |
| r(CAAU) | A-form | 33.4 | 49.9 |
| | Intercalated | 51.4 | 31.4 |
| | Bulge-out | 4.4 | 5.8 |
| r(GACC) | A-form | 73.5 | 86.6 |
| | Intercalated | 1.9 | - |
| | Loop | 7.0 | 2.2 |
| | Bulge-out | 5.5 | 1.7 |

[a] All simulations used 8 replicas and were run for 10 μs (the last 7 μs were used for data analysis). See Methods in the main text for details about clustering algorithm and Ref.[19] for details about structural states.

**Table S4.** Averaged values of global helical parameters (and their standard deviations) from 10 µs-long standard MD simulations of the r(GCACCGUUGG)$_2$ decamer excised from the 1QC0[20] structure and of the r(UUAUAUAUAUAUAA)$_2$ tetradecamer 1RNA.[21] [a]

| System/Setup | Helical rise [Å] | Helical incl. [°] | Tip [°] | Helical twist [°] | Major width [Å] | Minor width [Å] | X-disp. [Å] | Y-disp [Å] |
|---|---|---|---|---|---|---|---|---|
| 1RNA(Exp.) | 2.6 | 18.8 | -0.4 | 32.7 | 18.3 | 13.3 | -4.1 | -0.1 |
| OL3$_{CP}$–gHBfix19 | 2.5 ± 0.1 | 18.2 ± 0.8 | 0.0 ± 0.1 | 33.0 ± 1.1 | 19.2 ± 0.8 | 13.3 ± 0.4 | -4.2 ± 0.1 | 0.0 ± 0.0 |
| OL3$_{CP}$–NBfix19 | 2.5 ± 0.0 | 18.2 ± 0.5 | 0.0 ± 0.1 | 33.1 ± 0.2 | 19.2 ± 0.0 | 13.3 ± 0.0 | -4.1 ± 0.1 | 0.0 ± 0.0 |
| 1QC0 (Exp.) | 2.5 | 16.9 | 1.0 | 32.1 | 18.5 | 13.1 | -4.6 | 0.0 |
| OL3$_{CP}$–gHBfix19 | 2.7 ± 0.0 | 15.6 ± 0.6 | -0.5 ± 0.1 | 31.9 ± 0.2 | 19.2 ± 0.0 | 13.4 ± 0.0 | -4.7 ± 0.0 | 0.0 ± 0.0 |
| OL3$_{CP}$–NBfix19 | 2.7 ± 0.0 | 15.4 ± 0.6 | -0.4 ± 0.1 | 31.8 ± 0.2 | 19.2 ± 0.0 | 13.4 ± 0.0 | -4.7 ± 0.1 | 0.0 ± 0.3 |

[a] Simulation were run with the reference OL3$_{CP}$–gHBfix19 and OL3$_{CP}$–NBfix19 FFs (see Methods in the main text). The first two base pairs from each termini were excluded from the calculations to prevent end effects. Standard deviations were calculated using block average over 1000 snapshots (saved every 10 ps).

**Table S5.** Averaged values of base pair and local base pair step helical parameters (and their standard deviations) from 10 µs standard MD simulations of the r(GCACCGUUGG)$_2$ decamer excised from the 1QC0[20] structure and of the r(UUAUAUAUAUAUAA)$_2$ tetradecamer 1RNA.[21] [a]

| System/Setup | Shear [Å] | Stretch [Å] | Stagger [Å] | Buckle [°] | Propeller [°] | Opening [°] | Tilt [°] | Shift [Å] | Slide [°] | Rise [Å] | Roll [°] | Twist [°] | Fraying (top, bottom) [b] [%] | |
|---|---|---|---|---|---|---|---|---|---|---|---|---|---|---|
| 1RNA(Exp.) | -0.1 | -0.2 | 0.0 | 1.3 | -18.8 | -0.4 | 0.1 | 0.1 | -1.3 | 3.3 | 10.0 | 30.5 | - | - |
| OL3$_{CP}$–gHBfix19 | 0.0 ± 0.0 | 0.0 ± 0.0 | 0.1 ± 0.0 | 0.0 ± 0.2 | -17.4 ± 0.6 | 0.9 ± 0.2 | 0.0 ± 0.1 | 0.0 ± 0.0 | -1.5 ± 0.1 | 3.2 ± 0.1 | 10.5 ± 0.4 | 30.2 ± 1.0 | 3.9 | 5.4 |
| OL3$_{CP}$–NBfix19 | 0.0 ± 0.0 | 0.0 ± 0.0 | 0.1 ± 0.0 | 0.0 ± 0.2 | -17.4 ± 0.2 | 0.9 ± 0.2 | 0.0 ± 0.1 | 0.0 ± 0.0 | -1.5 ± 0.0 | 3.2 ± 0.0 | 10.5 ± 0.3 | 30.2 ± 0.1 | 4.9 | 4.6 |
| 1QC0 (Exp.) | 0.1 | -0.3 | 0.1 | 1.2 | -12.5 | 1.4 | -0.6 | 0.0 | -1.7 | 3.2 | 9.0 | 30.4 | - | - |
| OL3$_{CP}$–gHBfix19 | 0.0 ± 0.0 | 0.0 ± 0.0 | -0.1 ± 0.0 | 1.0 ± 0.2 | -12.4 ± 0.5 | -0.1 ± 0.1 | 0.2 ± 0.1 | 0.0 ± 0.0 | -1.8 ± 0.0 | 3.3 ± 0.0 | 8.6 ± 0.4 | 29.9 ± 0.2 | 0.1 | 0.3 |
| OL3$_{CP}$–NBfix19 | 0.0 ± 0.0 | 0.0 ± 0.0 | -0.1 ± 0.0 | 1.0 ± 0.2 | -12.3 ± 0.5 | 0.0 ± 0.1 | 0.2 ± 0.1 | 0.0 ± 0.0 | -1.8 ± 0.0 | 3.3 ± 0.0 | 8.5 ± 0.3 | 29.8 ± 0.3 | 0.1 | 0.2 |

[a] Simulation were run with the reference OL3$_{CP}$–gHBfix19 and OL3$_{CP}$–NBfix19 FFs (see Methods in the main text). The first two base pairs from each termini were excluded from the calculations to prevent end effects. Standard deviations were calculated using block average over 1000 snapshots (saved every 10 ps).
[b] Top and bottom base pairs are $G_1C_{20}$ and $G_{10}C_{11}$ for the 1QC0 duplex and $U_1A_{28}$ and $A_{14}U_{15}$ for the 1RNA duplex, respectively.

SUPPORTING FIGURES

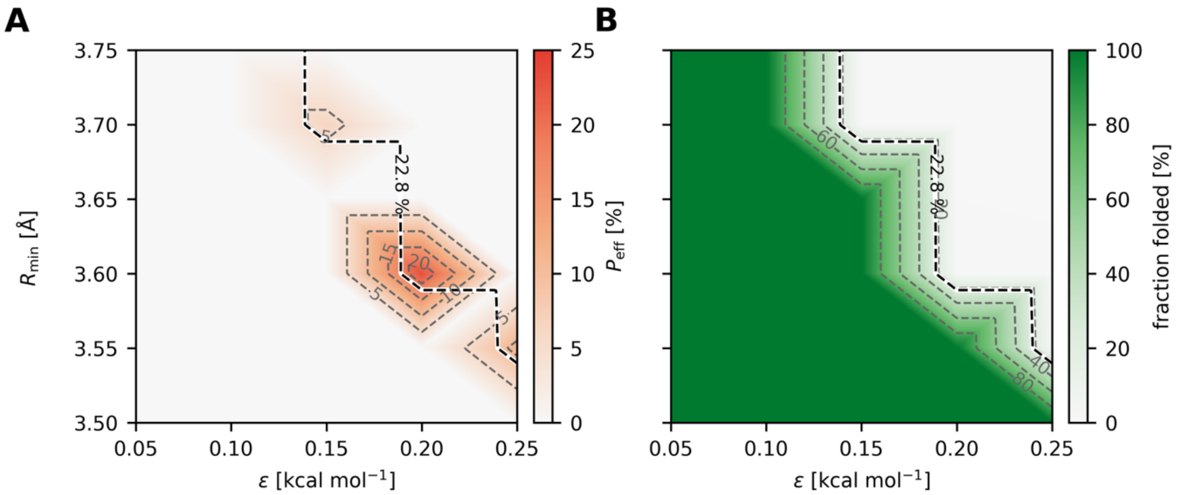


**Figure S1:** Reweighting-based optimization of NBfix parameters for the –N···N– interactions. (A) Normalized effective sample size ($P_{\mathrm{eff}}$) of the reweighted ensemble as a function of LJ radius *R* and well depth *ε*. (B) Reweighted folded-state fraction over the same range of parameters, with the dashed contour denoting the target folded-state population from the converged OL3$_{\mathrm{CP}}$–gHBfix19 T-REMD simulation.

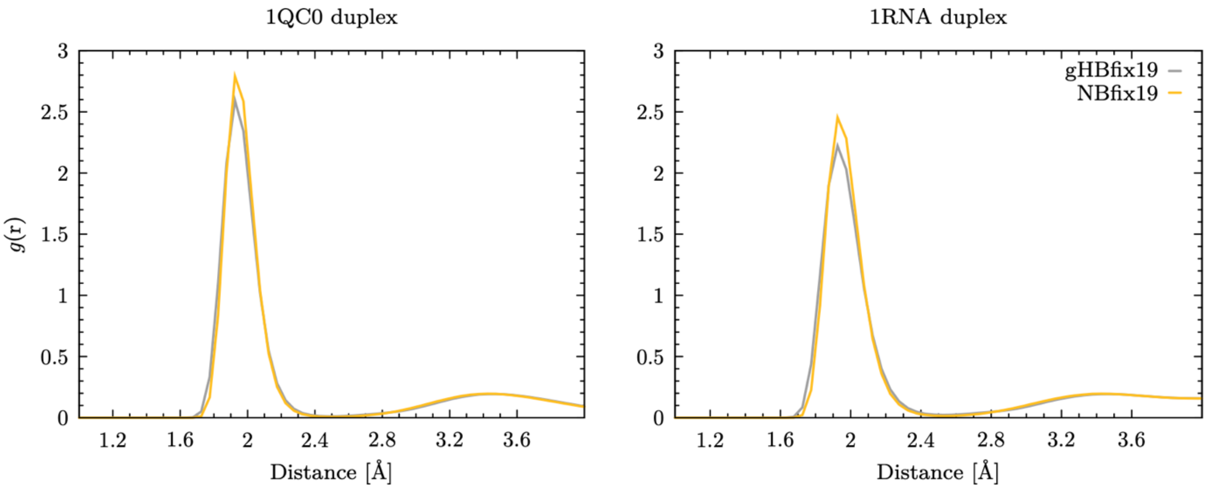


**Figure S2:** Radial distribution functions *g*(r) for –NH···N– interactions in the canonical RNA duplexes 1QC0 and 1RNA for the reference OL3$_{\mathrm{CP}}$–gHBfix19 and optimized OL3$_{\mathrm{CP}}$–NBfix19 FFs.

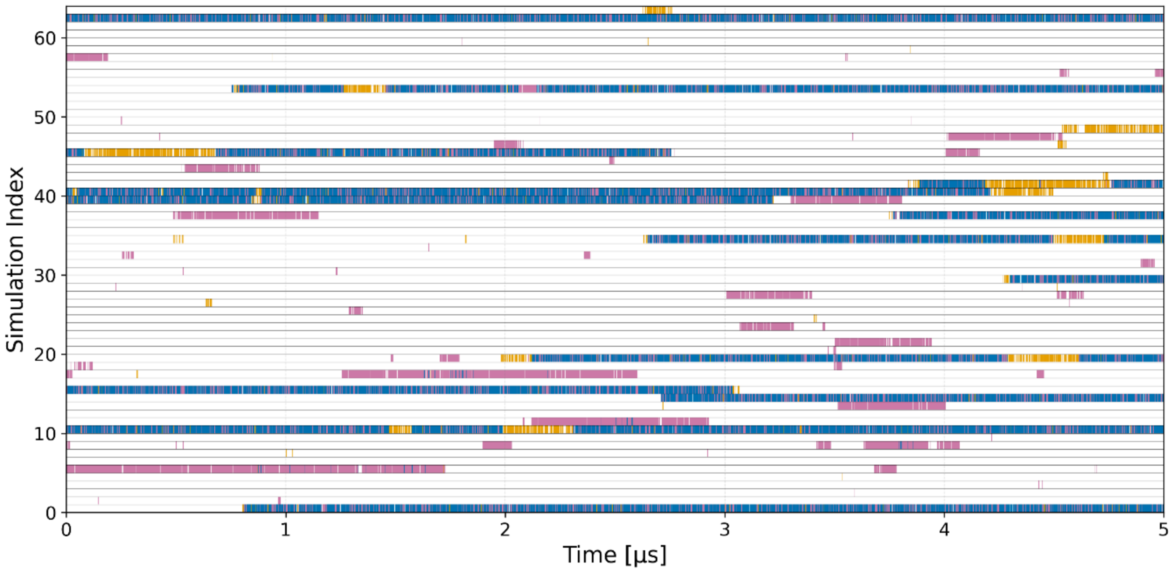


**Figure S3.** Conformational sampling of T-REMD folding simulation of the GAGA TL using the $OL3_{CP}$–gHBfix19–$NBfix_{NH...N}$ FF variant. The panel shows time evolution of major conformers, i.e., (i) correctly folded A-form stem and loop (native states with all signature interactions formed, blue), (ii) folded A-form stem (loop not in native conformation, purple), and (iii) correctly folded loop (stem not in A-form, orange), in all 64 continuous (demultiplexed) trajectories.

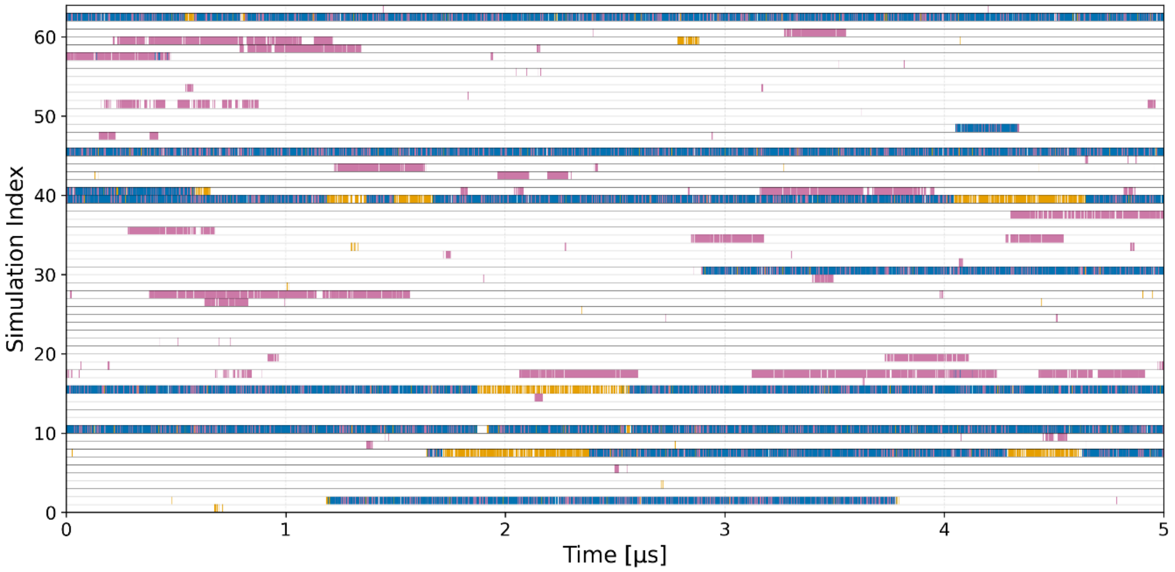


**Figure S4.** Conformational sampling of T-REMD folding simulation of the GAGA TL using the $OL3_{CP}$–gHBfix19–$NBfix_{OH...nbO}$ FF variant. See Figure S3 for more details.

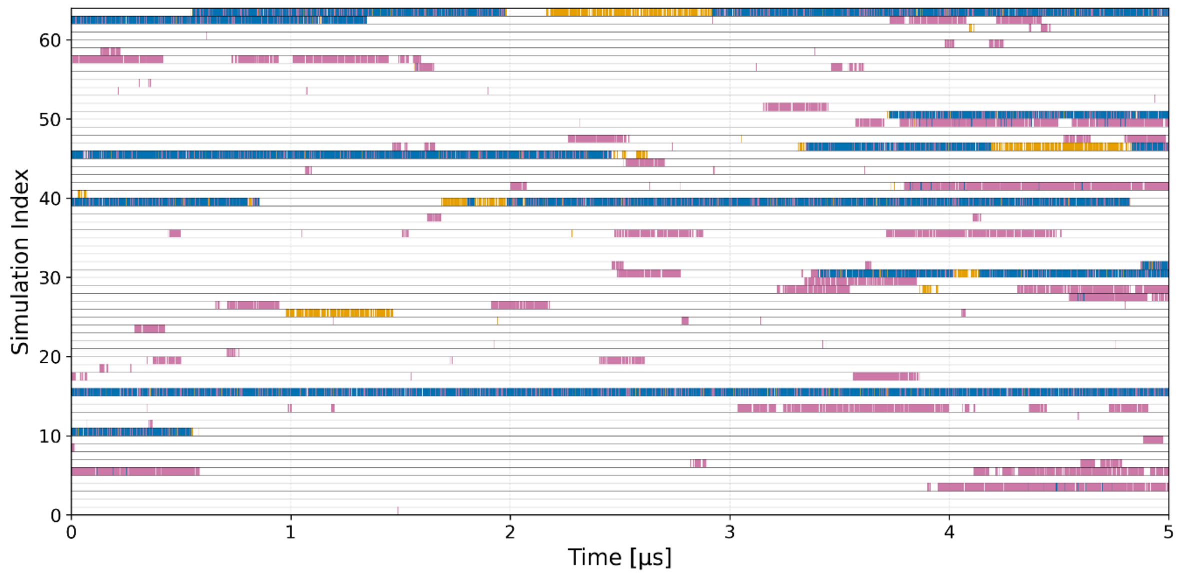


**Figure S5.** Conformational sampling of T-REMD folding simulation of the GAGA TL using the OL3$_{CP}$–gHBfix19–NBfix$_{OH...bO}$ FF variant. See Figure S3 for more details.

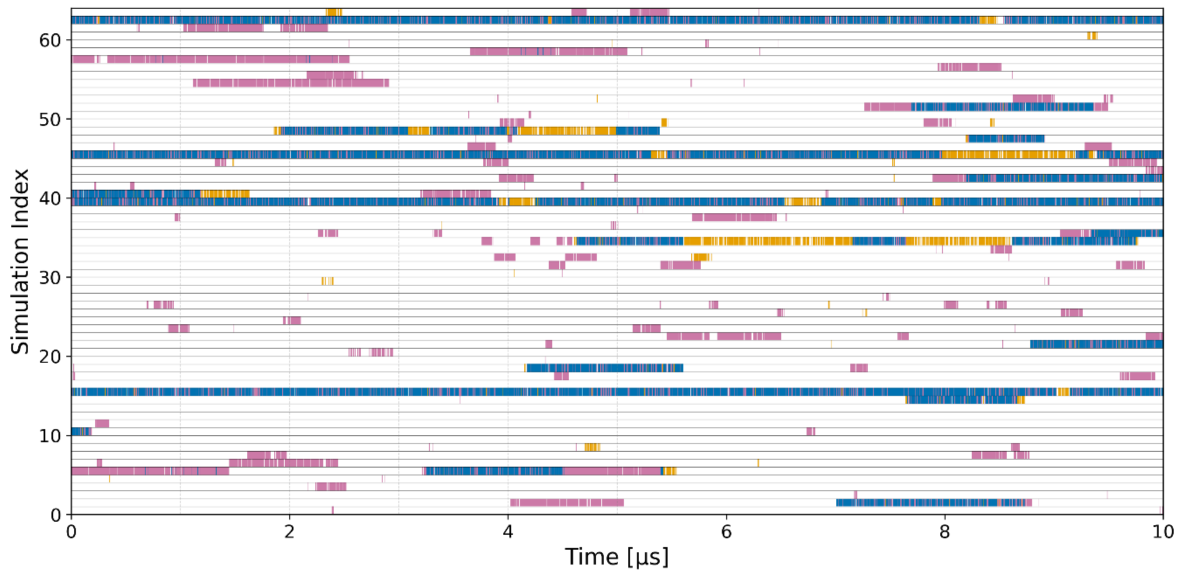


**Figure S6.** Conformational sampling of T-REMD folding simulation of the GAGA TL using the OL3$_{CP}$–NBfix19 FF. See Figure S3 for more details.

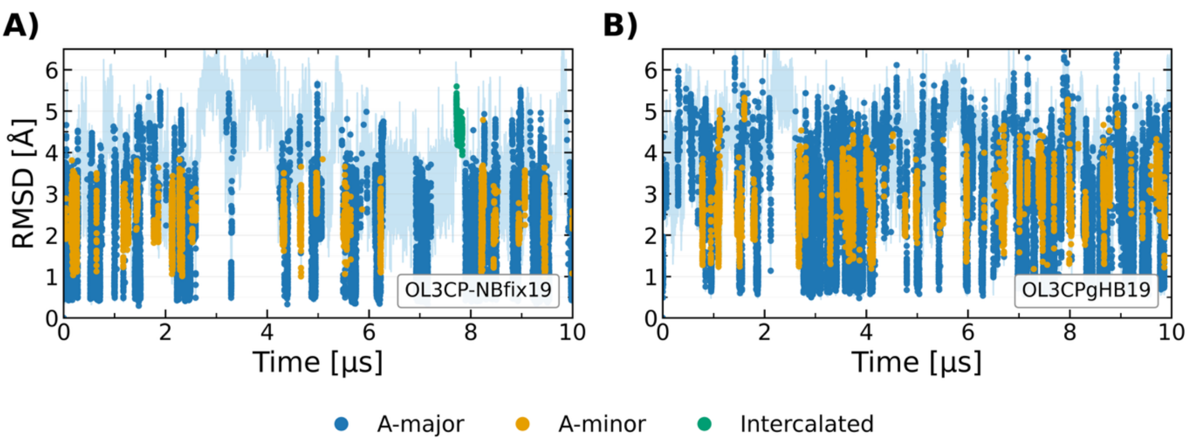


**Figure S7.** Time evolution of the RMSD with respect to the A-form conformation in two 10 µs-long standard MD simulations of the r(AAAA) TN using (A) $OL3_{CP}$–NBfix19 and (B) $OL3_{CP}$–gHBfix19 FFs. Light-blue traces show the RMSD time series. Colored markers denote frames assigned to reference conformational clusters based on the minimum εRMSD[22] to the cluster templates (assignment threshold εRMSD = 0.7): A-major (blue), A-minor (gold), and Intercalated (green). Frames that do not satisfy the assignment threshold are not classified and are therefore not shown.

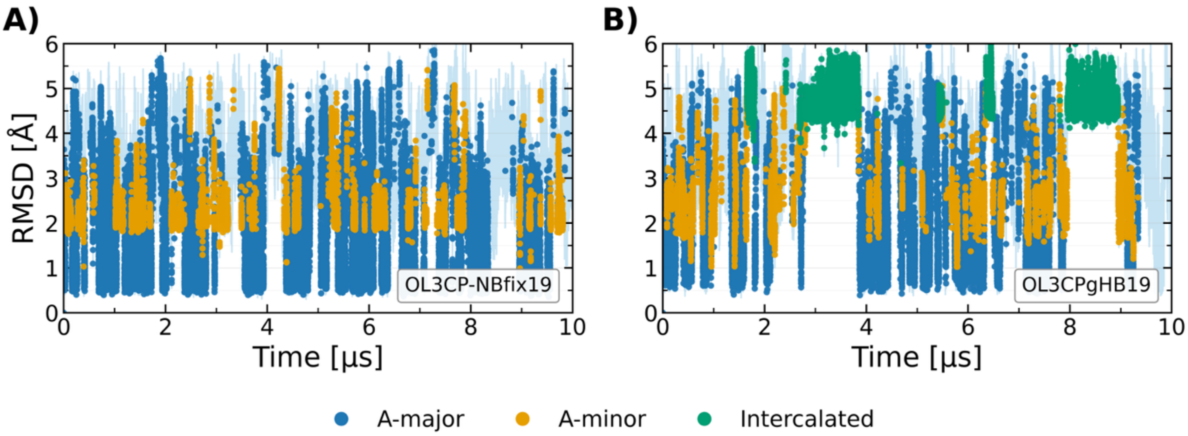


**Figure S8.** Time evolution of the RMSD in two 10 µs-long standard MD simulations of the r(CAAU) TN using (A) $OL3_{CP}$–NBfix19 and (B) $OL3_{CP}$–gHBfix19 FFs. See Figure S7 for more details.

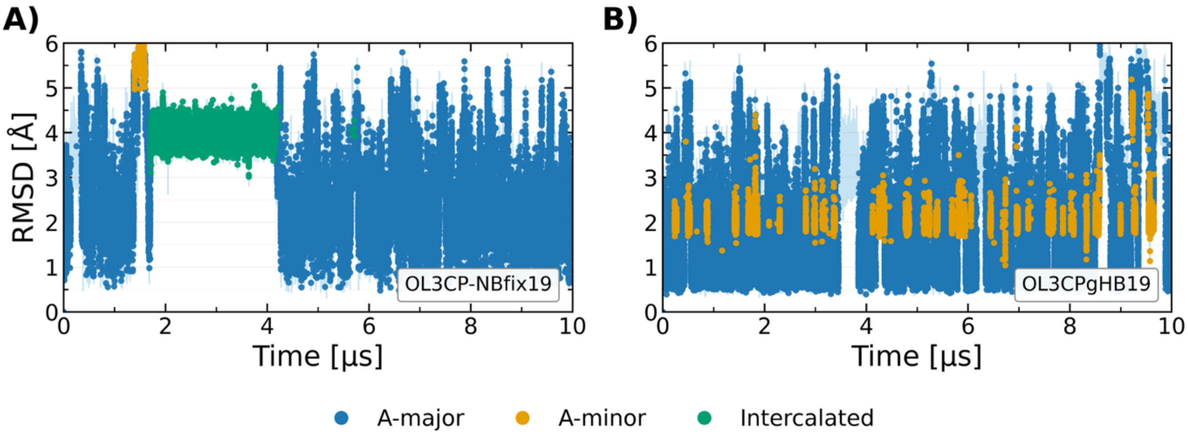


**Figure S9.** Time evolution of the RMSD in two 10 µs-long standard MD simulations of the r(CCCC) TN using (A) $OL3_{CP}$–NBfix19 and (B) $OL3_{CP}$–gHBfix19 FFs. See Figure S7 for more details.

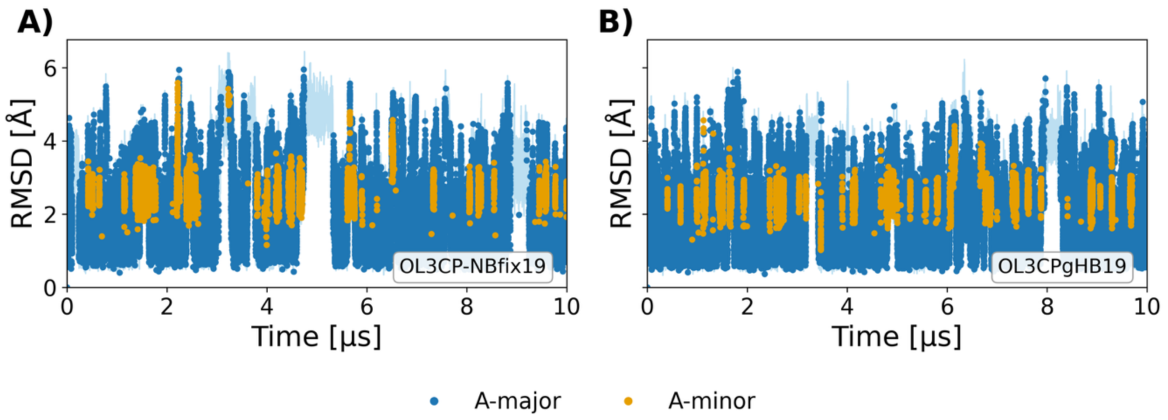


**Figure S10.** Time evolution of the RMSD in two 10 µs-long standard MD simulations of the r(GACC) TN using (A) OL3$_{CP}$–NBfix19 and (B) OL3$_{CP}$–gHBfix19 FFs. See Figure S7 for more details.

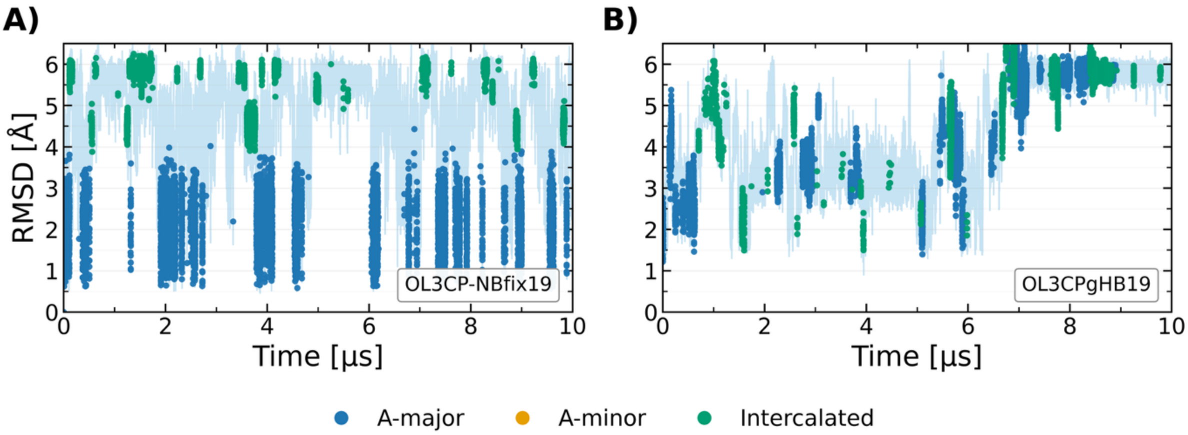


**Figure S11.** Time evolution of the RMSD in two 10 µs-long standard MD simulations of the r(UUUU) TN using (A) OL3$_{CP}$–NBfix19 and (B) OL3$_{CP}$–gHBfix19 FFs. See Figure S7 for more details.

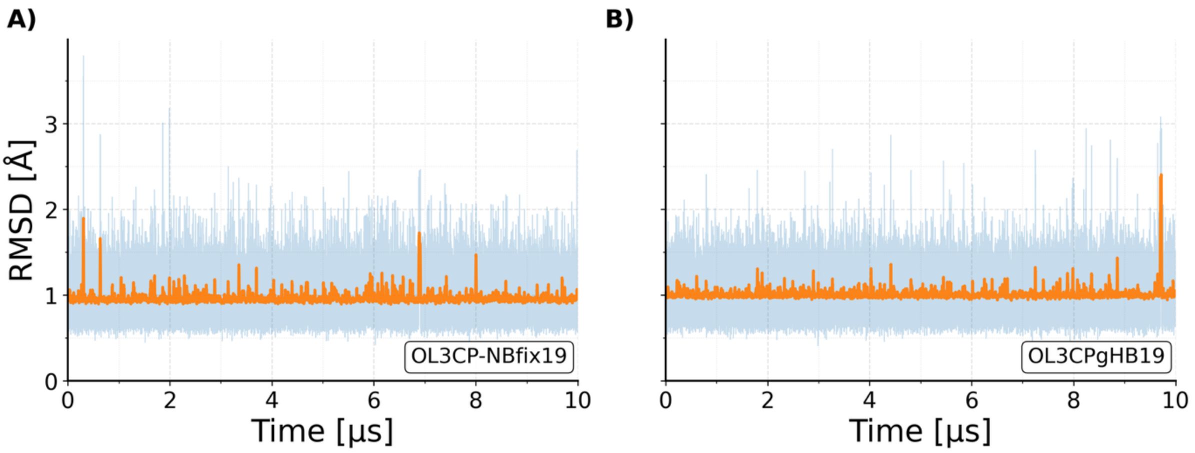


**Figure S12.** Evolution of the RMSD from two 10 µs-long standard MD simulations of the GAGA TL using (A) OL3$_{CP}$–NBfix19 and (B) OL3$_{CP}$–gHBfix19 FFs. Light blue traces show the actual RMSD fluctuations, whereas orange traces show the centered running average (10 ns windows).

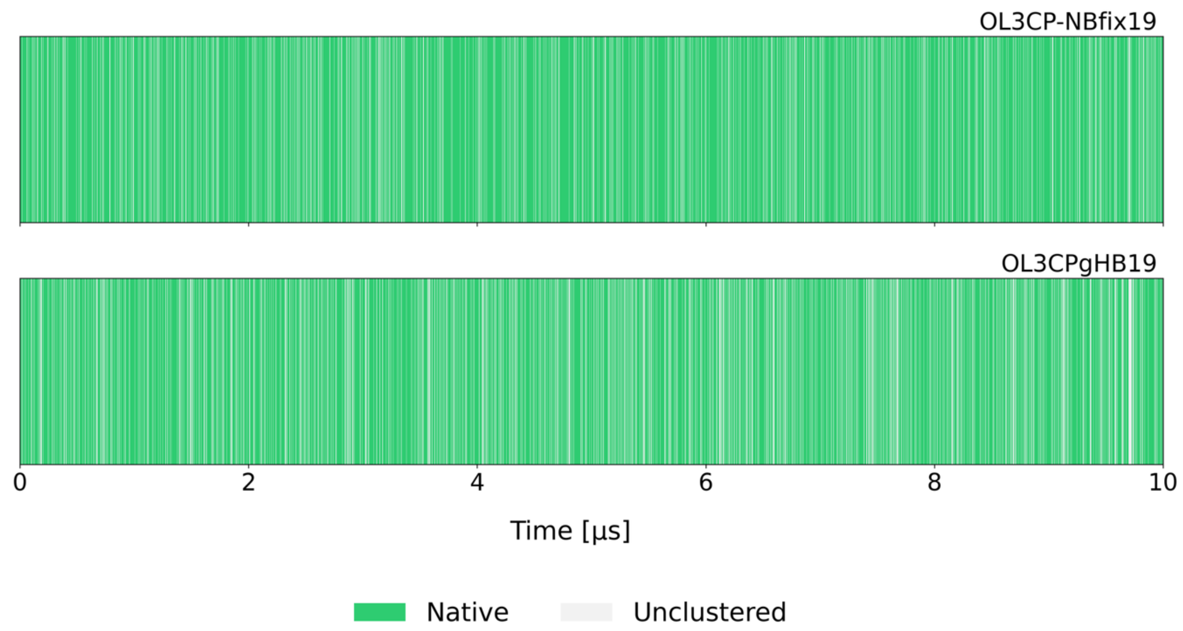


**Figure S13.** Time evolution of structural states sampled during two standard MD simulations of the GAGA TL using (A) OL3$_{CP}$–NBfix19 and (B) OL3$_{CP}$–gHBfix19 FFs. Each frame is classified as either native (green; based on the presence of all signature loop interactions) or unclustered (grey; representing conformations that do not satisfy the native-state criteria).

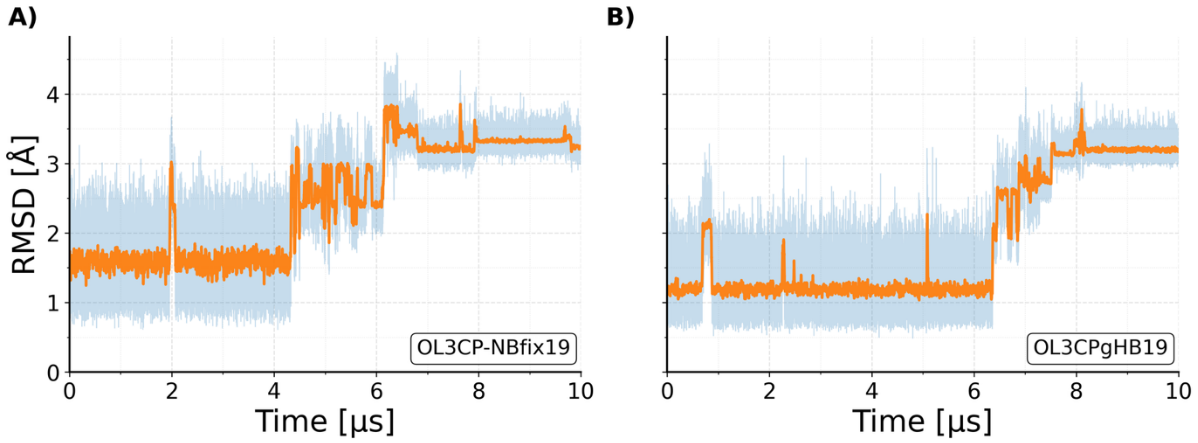


**Figure S14.** Evolution of the RMSD from two 10 µs-long standard MD simulations of the 14-mer UUCG TL (PDB ID 2KOC[8]) using (A) OL3$_{CP}$–NBfix19 and (B) OL3$_{CP}$–gHBfix19 FFs. See Figure S12 for more details.

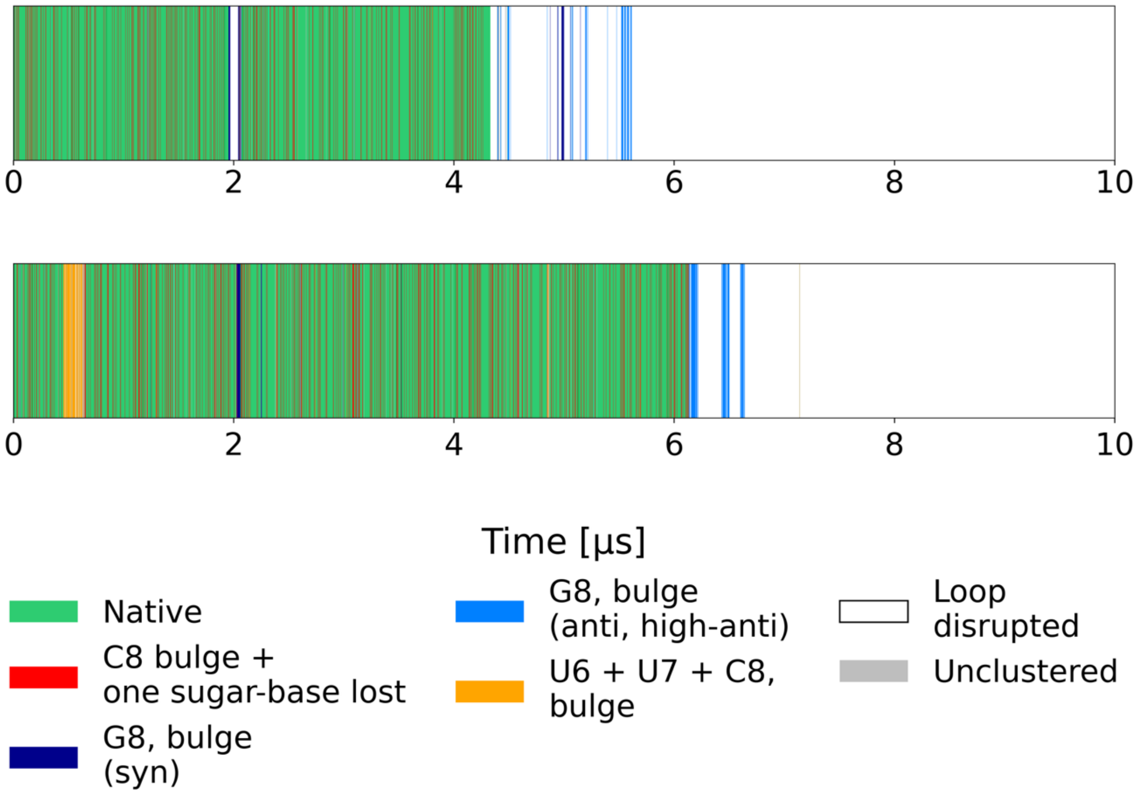


**Figure S15.** Time evolution of key structural states sampled during two standard MD simulations of 14-mer UUCG TL (PDB ID 2KOC[8]) using (A) $OL3_{CP}$–NBfix19 and (B) $OL3_{CP}$–gHBfix19 FFs. See Ref.[18] for details about the structural states.

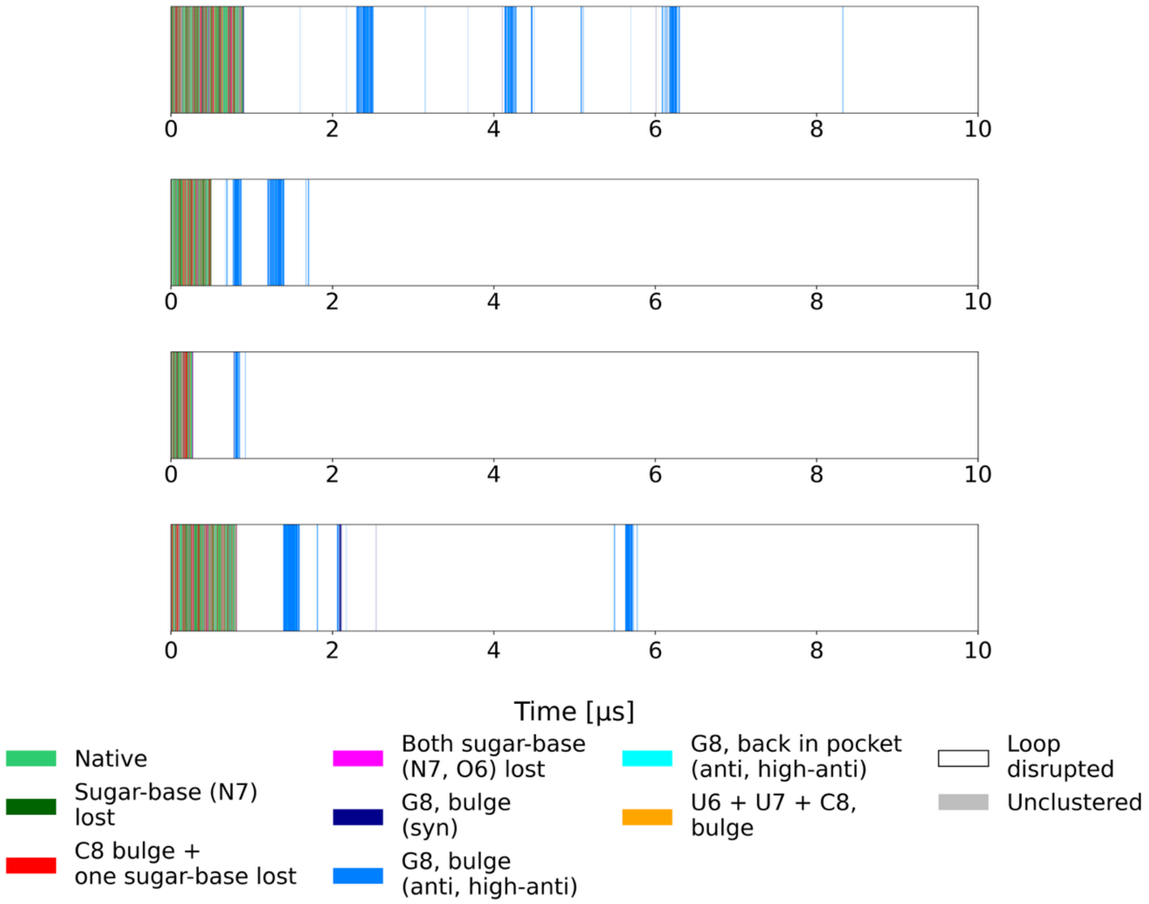


**Figure S16.** Time evolution of key structural states sampled during four independent standard MD simulations of 8-mer UUCG TL using OL3-vdW7-PAK FF. See Ref.[18] for details about the structural states.

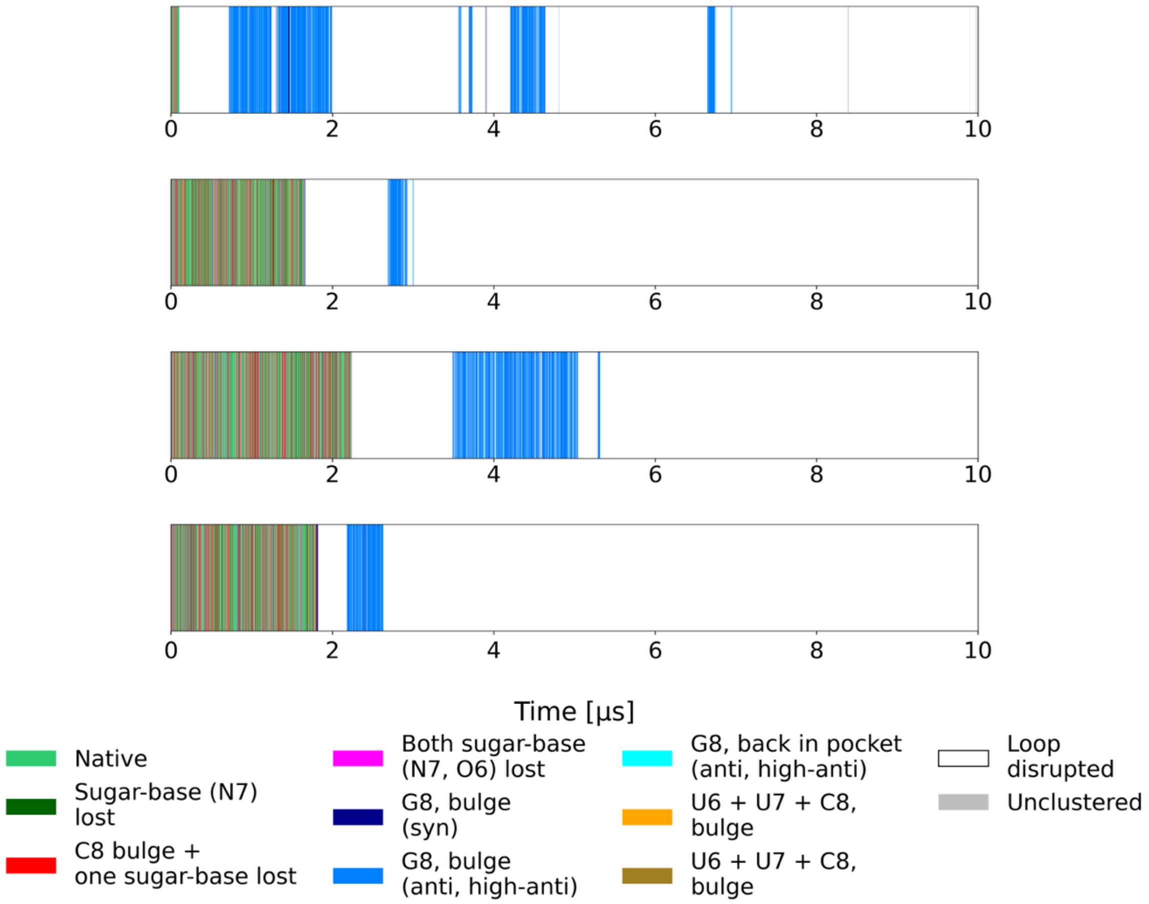


**Figure S17.** Time evolution of key structural states sampled during four independent standard MD simulations of 10-mer UUCG TL using OL3-vdW7-PAK FF. See Ref.[18] for details about the structural states.
